# A Conversational Multi-Agent AI System for Integrated Multi-Omics Analysis and Biomedical Discovery

**DOI:** 10.64898/2026.08.08.743577

**Authors:** Pankaj Rajdeo, Shunya Asanuma, Michal Kouril, Peixin Lu, Jichao Chen, Aman Chadha, V. B. Surya Prasath, Bruce J. Aronow, Nathan Salomonis

## Abstract

Single-cell and spatial omics offer unprecedented opportunities to decipher the mechanisms of disease, however, this process requires teams of experts, iterative trial-and-error and reasoning across modalities. Here we present LungChat (https://chat.lungmap.net), a conversational system for integrated multi-omics analysis and biomedical discovery, deployed as a hierarchical multi-agent architecture in which a supervisor decomposes natural-language questions into parallel, tool-grounded tasks spanning single-cell and spatial analyses, literature and clinical-trial synthesis, and drug repurposing. To predict new therapeutics, LungChat implements Direction-Aware Repurposing and Targeting (DART) to distinguish perturbations that reverse disease transcriptional programs from those that reinforce them, at the cell-type level, for safety prediction. Controlled architecture ablations showed that hierarchical orchestration improved grounded abstention and token efficiency and preserved strong performance on complex multi-step tasks. In pulmonary disease case studies, LungChat independently prioritized saracatinib for IPF through drug-connectivity screening, followed by DART-based cell-type analysis; the same compound has been evaluated in the STOP-IPF clinical trial (NCT04598919). The system also recovered fluticasone propionate, an established COPD therapy, through a single orchestrated analysis. This tissue-agnostic system provides a blueprint for verifiable agentic AI systems that support reproducible scientific discovery.

## 1 Introduction

Advances in single-cell and spatial transcriptomics have fundamentally changed how biological systems are studied, enabling cell-type–resolved and spatially contextualized analyses across development, homeostasis, and disease. Large-scale atlas efforts such as The Human BioMolecular Atlas Program (HuBMAP) [4], Human Cell Atlas (HCA) [21, 22], and LungMAP [6] have generated millions of profiles across the spectrum of human tissues, development and disease, creating unprecedented opportunities for systems-level discovery. However, the increased scale and complexity of these datasets, shifts the challenge from analysis to systems-level integration. This challenge necessitates integration of insights across omics technologies, while reconciling inconsistent metadata across studies.

Existing solutions address parts of this problem but remain structurally limited. Interactive visualization platforms such as the CZI CellxGene, and UCSC cell browsers [20, 26] enable exploration of atlas-level datasets but lack support for deep, quantitative analysis. General-purpose AI assistants (e.g., ChatGPT, Claude, etc) can synthesize vast amounts of prior knowledge from the literature but lack harmonized quantitative data, metadata or access to validated analysis tools, resulting in "best guesses" [12]. As a result, there is a growing disconnect between the availability of atlas-scale data and the ability of researchers to interrogate it rigorously and iteratively.

Recent progress in large language models (LLMs) and agentic AI systems suggests a path forward. Conversational interfaces have demonstrated promise for lowering the barrier to complex analyses [23]. Pre-trained language models and single-agent approaches, such as CellWhisperer [23], CompBioAgent [29], Cell2Text [13] and scChat [17] generate rapid and automated annotation and visualization of single-cell genomics data. Multi-agent architectures such as BioMaster [27], BioAgents [18] and Biomni [10] enable inter-agent coordination, planning and task execution to nominate hypotheses for validation. Existing approaches such as DrugAgent [11] and TxGNN [9], provide exciting new approaches to augment expert-driven drug repositioning. Despite these innovations, numerous challenges remain in AI assisted drug discovery and biomedical reasoning, including standardized integration of findings across studies and modalities (space, time, and disease); functional prediction (pathways, regulatory potential, and cellular interactions); assessment of the potential positive and negative impacts of new therapies; and contrasting findings across disease and developmental contexts.

In this work, we introduce LungChat, a conversational multi-agent AI system for integrated multi-omics analysis, designed to support transparent, reproducible biomedical discovery through natural language interactions. LungChat is open-access at https://chat.lungmap.net and leverages 39 single-cell datasets spanning 2.8 million cells, integrated spatial omics technologies and over 50 analytical tools. The system combines (i) **multi-agent orchestration**, in which a supervisor coordinates specialized sub-agents; (ii) **tool-grounded analysis**, grounding language model outputs in structured execution of validated bioinformatics methods including functional enrichment (ToppGene [5]), drug connectivity and repurposing (iLINCS [19]), literature retrieval and synthesis (PubMed); and (iii) **provenance-tracked reproducibility**, where every analytical step produces persistent, publication-ready outputs with complete parameter provenance. For therapeutic hypothesis generation, we developed Direction-Aware Repurposing and Targeting (DART), a method for cell-type-resolved drug prioritization that classifies perturbations as repression or activation to predict on-target efficacy and off-target safety at cell-type resolution (Section 4.7). Applied to idiopathic pulmonary fibrosis (IPF), LungChat independently prioritized saracatinib through drug-connectivity screening, followed by DART-based cell-type analysis; the same compound has been evaluated in the STOP-IPF clinical trial (NCT04598919). In Chronic Obstructive Pulmonary Disease (COPD), which affects over 300 million individuals, DART identifies fluticasone propionate, a corticosteroid used in COPD combination therapy, among the top predicted disease-reversal signatures. By aligning natural language interaction with real analytical execution, this work positions conversational multi-agent systems as a practical interface for atlas-scale data exploration. The system complements emerging AI scientist approaches [7, 16] by prioritizing transparent, tool-grounded analysis over autonomous hypothesis generation, addressing a critical gap between accessibility and rigor in contemporary biomedical research.

## 2 Results

### 2.1 Coordinated multi-agent analysis across omics and biomedical knowledge

LungChat was developed to translate complex biomedical questions into structured analyses by leveraging harmonized omics datasets, reproducible analytical results, publication-ready visualizations, and literature grounded evidence synthesis. Where possible, computationally intensive analyses are precomputed, enabling rapid retrieval of results from harmonized datasets and metadata. Each analytical step is grounded in the structured execution of validated bioinformatics methods and produces persistent, publication-ready outputs with complete parameter provenance, allowing analyses to be inspected, reproduced, and extended. Rather than generating plausible but unverified responses, LungChat returns evidence-grounded results that researchers can examine and rerun.

LungChat implements a hierarchical multi-agent architecture with a central stateful supervisor agent that coordinates the analysis and maintains the conversational state that serves as the system’s working memory (Figure 1). This statefulness enables multi-turn, iterative dialogue in natural language. The supervisor coordinates a team of specialized subagents, each augmented with a library of validated analytical tools: a single-cell agent for differential expression, cell-type characterization, correlation, and visualization; a spatial agent for spatial gene-expression mapping, cell-cell communication inference, and neighborhood enrichment within and across technologies; a research agent that bridges internal data with external knowledge through functional enrichment, literature synthesis, clinical-trial retrieval, drug connectivity, and repurposing; and a LungMAP agent for natural-language exploration of the LungMAP consortium database. For analyses not covered by the core tools, the supervisor can generate and execute custom code in a secure sandbox. Functional enrichment is performed with ToppGene, drug connectivity and repurposing through iLINCS, and literature synthesis across EuropePMC. User queries are decomposed into tractable subtasks that execute in parallel when independent, and their results are synthesized into coherent scientific responses. To increase analytical throughput, the supervisor can invoke multiple subagents concurrently, including multiple instances of the same subagent when appropriate.

**Figure 1:**
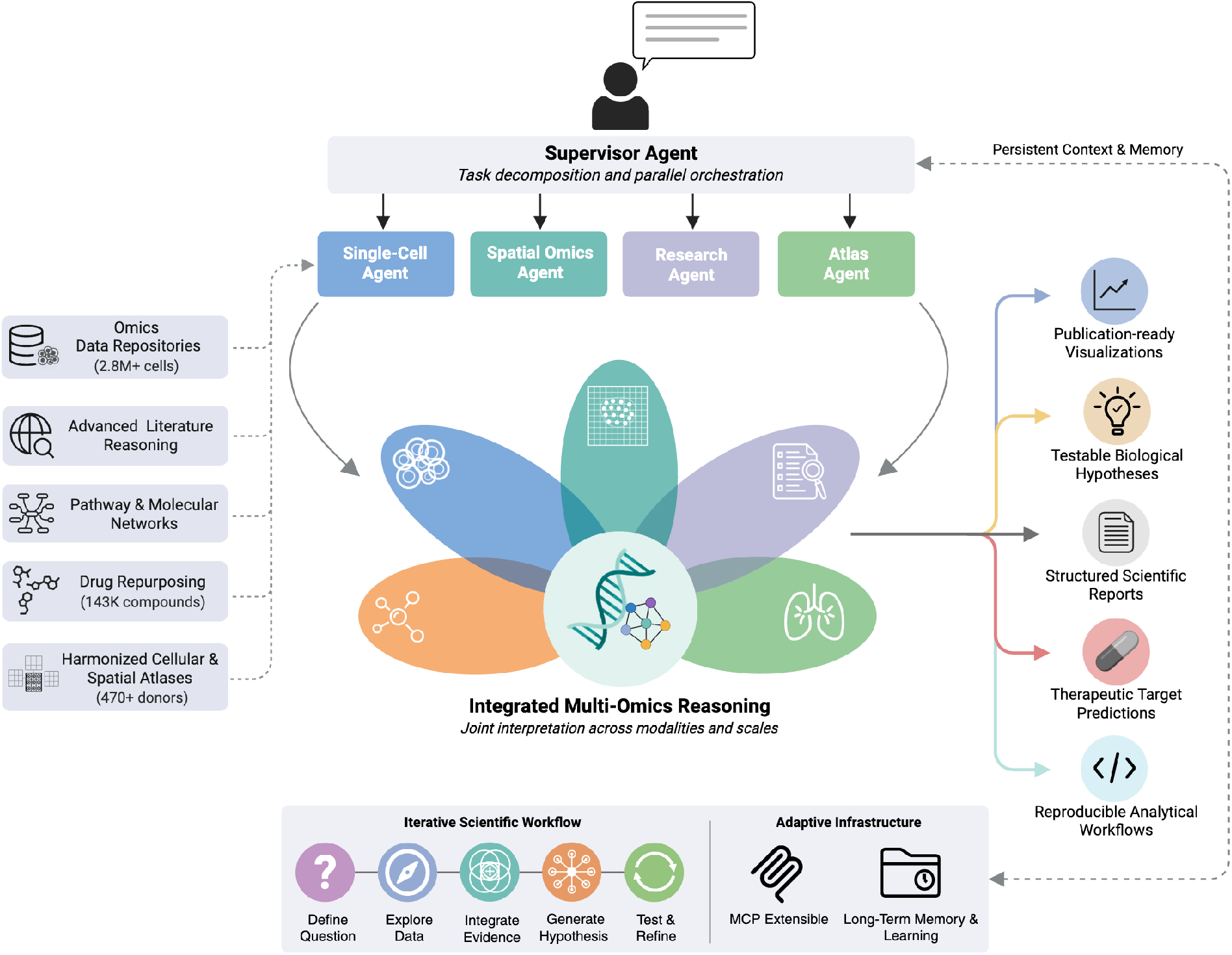
LungChat system architecture and workflow. A hierarchical multi-agent system integrating conversational AI with multi-omics analysis. The Supervisor Agent orchestrates four specialized subagents (Single-Cell, Spatial, Research, Atlas) that access curated data repositories (scRNA-Seq, spatial transcriptomics, drug compounds, regulatory networks, literature) to generate publication-ready outputs including visualizations, testable hypotheses, therapeutic predictions, and reproducible workflows. The system supports iterative scientific workflows (Define → Explore → Integrate → Generate → Test & Refine) with long-term memory and MCP-extensible infrastructure for adaptive tool integration.

For therapeutic hypothesis generation, the research agent’s drug connectivity and repurposing analysis is coupled to Direction-Aware Repurposing and Targeting (DART), a method for cell-type-resolved drug prioritization that classifies perturbations as reversing or reinforcing a disease transcriptional program to predict on-target efficacy and off-target safety at cell-type resolution.

### 2.2 Benchmarking and architecture ablation

We evaluated LungChat using a comprehensive benchmark comprising 100 test cases across five evaluation tracks: 50 single-step analytical queries (analytical workflows, data visualization, metadata and knowledge queries), 13 security and adversarial queries (prompt injection, security exploitation, out-of-domain), 17 grounded abstention queries testing appropriate scope limitation, 10 multi-tool orchestration queries requiring cross-domain coordination, and 10 decomposition-stress queries testing scientific reasoning depth. To assess the contribution of the multi-agent architecture, we conducted ablation experiments across four configurations. Following current evaluation best practices [3], we employed both deterministic code-based evaluators (configuration correctness, execution success, metadata consistency, tool selection accuracy) and LLM-as-judge scoring using GPT-4o (temperature=0) with a structured binary rubric covering biological accuracy, visual correctness, and refusal appropriateness. Each test case was executed across three independent trials (*n*=3 repetitions per case) to assess reliability.

To isolate the contribution of hierarchical multi-agent orchestration, we evaluated LungChat under four ablation configurations: (1) **MA+Sel** (production), the hierarchical architecture with hybrid tool selection; (2) MA−Sel, the production hierarchy without the tool selector; (3) Flat+Sel, a single agent with all 51 tools and merged domain prompts, retaining the tool selector; and (4) **Flat**−**Sel**, a single agent with all tools, merged prompts, and no selector. Flat-agent baselines preserve all domain knowledge from specialist prompts, ensuring differences reflect architectural design rather than information asymmetry.

Figure 2 summarizes the ablation results across all five tracks (100 questions, 1,080 total evaluations). On single-step queries (Figure 2a), all four configurations achieved ≥89% LLM Judge accuracy, with Flat−Sel scoring highest (98.0%), confirming that single-step quality is architecture-neutral. The hierarchical architecture achieves a consistent ∼2× reduction in token use on single-step queries and other lower-complexity evaluation tracks (Figure 2b), reflecting the context-partitioning benefit of scoped tool pools described in Section 4.4.2. Grounded abstention (Figure 2c) revealed the largest architectural difference: MA+Sel blocked 90.2% of disallowed tool invocations, compared with 54.9% for Flat−Sel, a difference of 35.3 percentage points. On complex multi-tool and decomposition-stress queries (Figure 2d), both architectures achieve comparable quality, confirming that analytical depth stems from domain engineering rather than hierarchy. Full per-track results are provided in Appendix F.

**Figure 2:**
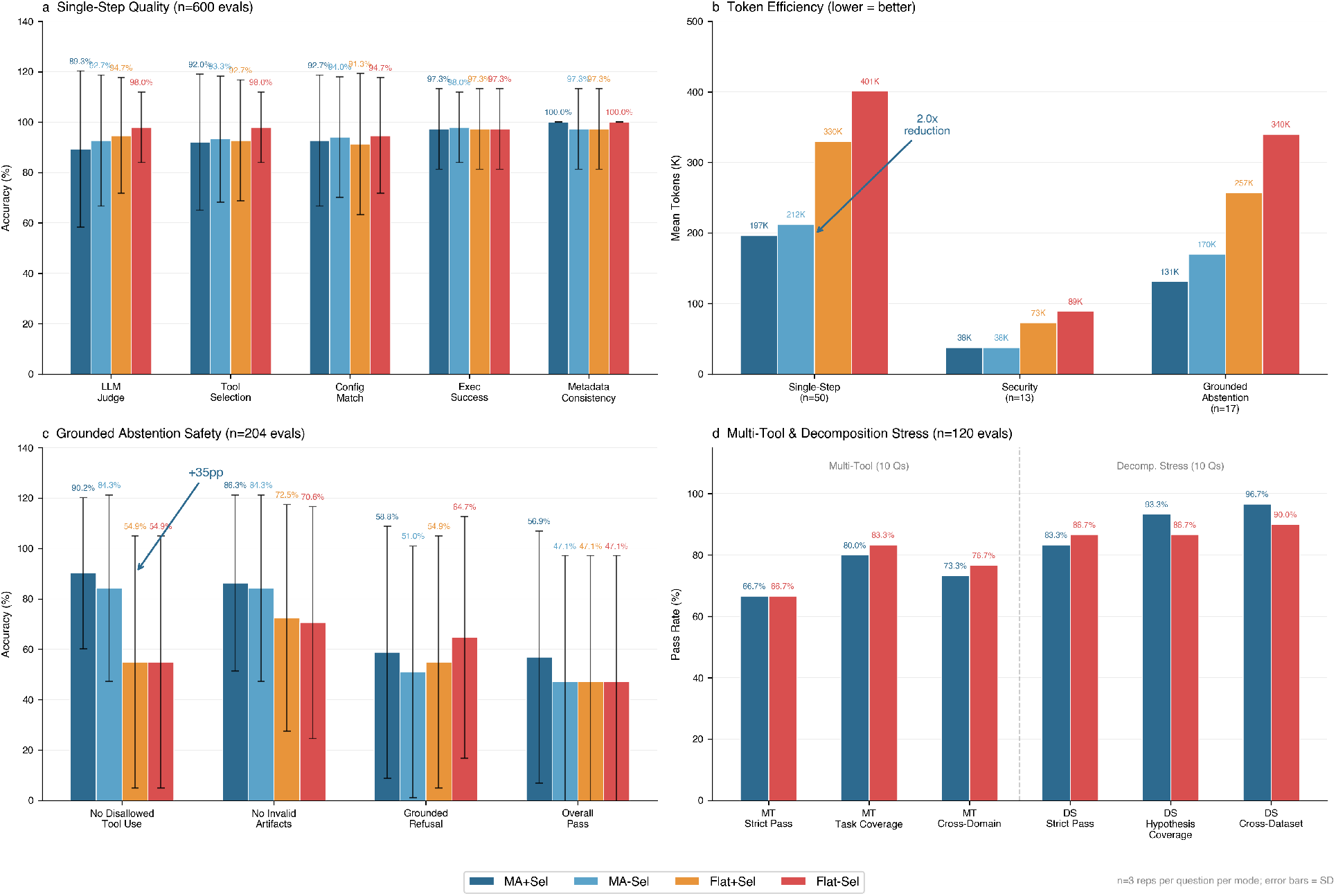
Architecture ablation evaluation across all five tracks (1,080 evaluations). (a) Single-step quality across four ablation configurations (50 queries × 3 reps × 4 modes = 600 evaluations), (b) mean token consumption per query across three evaluation tracks (lower is better), (c) grounded abstention safety metrics (17 queries × 3 reps × 4 modes = 204 evaluations), showing the +35pp improvement in disallowed-tool blocking for the multi-agent configuration, and (d) multi-tool orchestration and decomposition-stress results (20 queries × 3 reps × 2 modes = 120 evaluations), demonstrating architecture-neutral quality on complex tasks.

Across the tested adversarial tasks, the system prevented all prompt-injection, developer-mode, and instruction-override attempts, as well as all attempts to perform system reconnaissance or extract environment variables. All analytical outputs are saved with complete parameter provenance to support reproducible re-execution (Appendix C).

### 2.3 Cell-type-resolved therapeutic prioritization and mechanistic exploration

To evaluate LungChat’s ability to plan and execute multi-step workflows and synthesize their results, we focused on pulmonary fibrosis, for which effective therapeutic options remain limited. Existing therapies have largely aimed to suppress the activation, migration, and proliferation of resident lung fibroblasts, processes that contribute to tissue scarring and impaired lung function.

As a first evaluation, we explicitly asked LungChat to identify therapeutic targets for pulmonary fibrosis based on differential expression, pathway analysis, spatial location, and cell communication analyses. Our supervisor agent decomposed this query into six coordinated subtasks, executing independent analyses in parallel and dependent analyses sequentially (Figure 3a,b).

**Figure 3:**
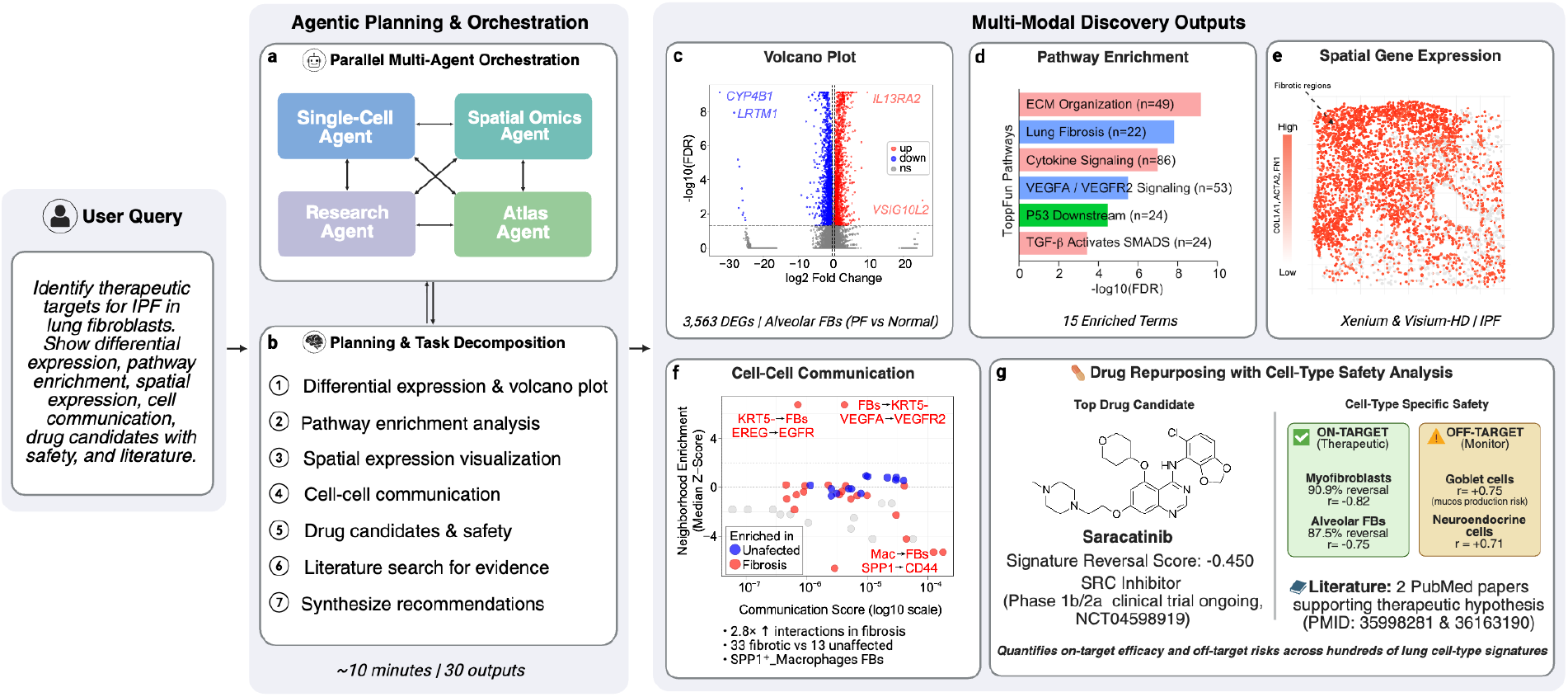
Automated therapeutic target and drug prioritization for pulmonary fibrosis. (a) LungChat parallel multi-agent orchestration with supervisor-coordinated task delegation. (b) Planning and task decomposition for reproducible workflows. (c) Ranking of up- and down-regulated genes in pulmonary fibrosis alveolar fibroblasts. (d) Gene-set enrichment (ToppGene) identifying known druggable pathways in pulmonary fibrosis. (e) Spatial transcriptomic localization of fibrotic markers. (f) Prediction of receptor-ligand interactions ranked by cell-type spatial proximity in pulmonary fibrosis versus controls (Xenium) cell-cell communication strength. (g) Drug repurposing with DART cell-type safety analysis: Saracatinib identified as top candidate. Off-target monitoring identified goblet cells and neuroendocrine cells.

**Figure 4:**
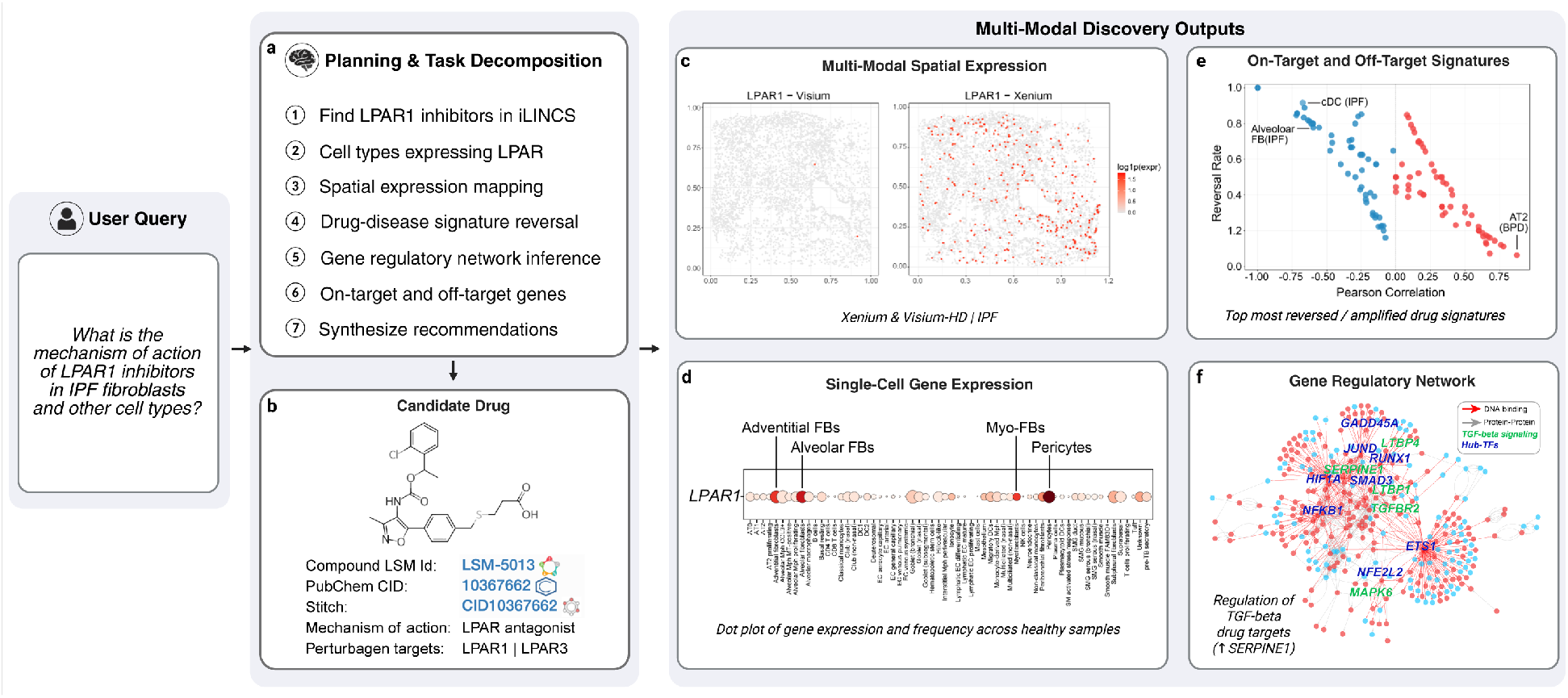
Resolving the regulatory mechanisms and cell targets of new therapies. (a) Planning and task decomposition to define diverse mechanisms of action for LPAR1 inhibitors in pulmonary fibrosis. (b) Identification of LPAR1 inhibitors in iLINCS with in vitro gene expression responses. (c) Comparison of the spatial distribution LPAR1 in Xenium and Visium-HD (same slide - Spatial Agent). (d) Expression and frequency of LPAR1 across adult healthy lung cell populations (HLCA - Single Cell Agent). (e) DART prediction of the top impacted lung disease, technology and developmental perturbations (Omics Data Repository), reversed or further amplified by in vitro LPAR agonist treatment (iLINCS). (f) Inferred gene regulatory network from annotated transcription factor (TF) target and pathway interactions (NetPerspective), with highlighted major TF nodes and annotated TGF-beta signaling responsive genes (ToppGene).

These analyses successfully identify well-defined pulmonary fibrosis marker genes (e.g., IL13RA2, COL1A1) and enriched pathways of IPF (e.g., extracellular matrix deposition, P53 transcriptional regulation, VEGFA and TGF-*β* signaling; Figure 3c-e). Integrated spatial-based cell-communication analyses (Squidpy, CellChat) further nominated EREG and VEGFA signaling from aberrant basaloid cells (pulmonary fibrosis KRT5-/KRT17+ alveolar type 2 cells) to activated fibroblasts, as uniquely spatial enriched interactions in pulmonary fibrosis versus controls (Figure 3f). This result is supported by VEGFA/VEGFR2 signaling from unbiased global gene set enrichment (Figure 3d). LungChat performed drug repurposing analysis via iLINCS connectivity mapping combined with DART cell-type safety profiling. In addition to recovering the approved IPF therapy nintedanib and the drug-associated target PDGFRB, LungChat identified saracatinib, a selective SRC kinase inhibitor, as the top therapeutic candidate (Figure 3g). DART cell-type impact analysis across 475 lung cellular signatures demonstrated that Saracatinib exhibits 90.9% reversal of pathogenic myofibroblast signatures (Pearson *r* = −0.82) and 87.5% reversal of alveolar fibroblast activation (*r* = −0.75), with an average fibroblast specificity of 54.8%. Off-target safety profiling identified potential risks in fetal/adult smooth muscle and pericytes, affecting approximately a dozen genes in each. Saracatinib had previously demonstrated superior anti-fibrotic efficacy compared to nintedanib and pirfenidone in preclinical models [1] and has been evaluated in STOP-IPF (NCT04598919), providing external support for the overall prioritization. This therapeutic target identification was performed as a single natural-language prompt submitted to a fresh LungChat session with no prior conversational context or iterative prompt refinement; the complete execution trace, including all tool calls, parameters, and intermediate outputs, is provided in Appendix E.

As a separate evaluation, we asked LungChat to broadly predict drugs that would reverse disease gene expression profiles in any (separate) or all (union) cell populations from patients with COPD. This analysis identified fluticasone propionate, an inhaled corticosteroid used in COPD combination therapy, as among the top predicted anti-COPD drug reversal signatures in dendritic cells. Fluticasone Propionate is a synthetic trifluorinated corticosteroid which reduces airway inflammation in moderate-to-severe COPD in concert with *β*2-agonists, such as Fenoterol. Fenoterol was identified among the top 3 anti-COPD drug reversal signatures, when all cell type COPD DEGs were aggregated (union) by LungChat. DART predicted off-target reversal of fetal and adult normal gene expression profiles in fetal smooth muscle, neonatal secondary crest myofibroblasts and adult ionocytes. Given the rarity of ionocytes and relative recent nature of their functional characterization, such cellular impacts would likely go unnoticed in conventional off-target screening cellular platforms.

#### 2.3.1 Regulatory mechanism evaluation

In addition to the discovery and safety evaluation of promising therapeutic candidates, LungChat can deeply evaluate regulatory mechanisms from a diverse repertoire of omics-focused analyses and the scientific literature. To assess this capability, LungChat was tasked to evaluate the potential mechanisms of action of a new drug target under current FDA evaluation. Lysophosphatidic acid receptor (LPAR) inhibitors target TGF-beta signaling pathway members, which are well-documented to be up-regulated in pulmonary fibrosis. To determine the mechanism of action of these inhibitors, LungChat executed a multi-step analysis to: 1) identify candidate drugs in iLINCS, 2) determine which cell types express LPAR1, 3) identify its spatial niches, 4) determine which cell-type disease programs could be reversed by drug treatment and to identify which transcriptional regulators were upstream of pharmacologically reversed genes (transcriptionally). These analyses confirmed expression of LPAR1 in all adult lung fibroblast and pericyte populations, confirmed spatial expression of LPAR1 with activated fibroblasts in the IPF niche, identified expected and novel IPF cellular impacts (reversal of IPF expression in fibroblasts and myeloid cells, respectively) and identified SMAD3 as the upstream transcriptional regulator of LPAR inhibitor targets genes (upregulation of SERPINE1). However, DART identified approximately a dozen potential off-target effects, including in presumably healthy smooth muscle, lymphatic endothelial, submucosal gland, deutrosomal, fetal Schwann cell and myofibroblasts with LPAR1 inhibition. Thus, LungChat is able to autonomously define complex regulatory mechanisms underlying drug treatment, including known and novel predicted interactions.

**Table 1:** LungChat data and system coverage.

| Resource | Coverage |
| --- | --- |
| Single-cell datasets | 39 (2.8M cells) |
| Spatial datasets | 1 (Visium-HD, Xenium) |
| Disease contexts | 15 |
| Specialized tools | 51 |
| Precomputed DEG signatures | 579 |

## 3 Discussion

We present LungChat, a conversational multi-agent AI system for integrated multi-omics analysis that supports transparent, end-to-end scientific workflows across single-cell and spatial transcriptomic data, along with pseudobulk expression summaries, while reducing the need for manual scripting. Architecture ablation across five evaluation tracks (1,080 evaluations) showed that the evaluated architectures achieved broadly comparable analytical quality on complex tasks, while the multi-agent hierarchy provided approximately 2× greater token efficiency and a 35-percentage-point improvement in grounded abstention. These results support the value of context partitioning and hierarchical orchestration for efficient and appropriately constrained analytical systems.

Through a series of therapeutically focused case studies, we demonstrate that LungChat can recover established disease-associated biology, characterize regulatory processes, compare diseased and healthy spatial niches, and identify known antifibrotic therapies. Beyond recapitulating known biology, we introduce Direction-Aware Repurposing and Targeting (DART), a cell-type-resolved method that assesses whether perturbation-induced transcriptional signatures reverse or reinforce disease-associated programs. Applied to IPF, LungChat independently prioritized saracatinib through drug-connectivity screening, followed by DART-based cell-type analysis; the same compound has been evaluated in the STOP-IPF clinical trial (NCT04598919), providing external support for the biological relevance of this prioritization. In COPD, DART recovered fluticasone propionate, an established therapy, supporting the ability of the method to identify clinically relevant perturbations from cell-type-resolved disease signatures.

More broadly, DART can compare disease-associated, developmental, and perturbation-induced transcriptional programs across cell types, datasets, and technologies. Such comparisons may reveal conserved biological programs, technology-associated differences, and potentially analogous or inconsistently annotated cell populations across studies. These analyses could support drug-discovery workflows by identifying molecular pathways, cell populations, and disease contexts predicted to be reversed or reinforced by a perturbation. By revealing heterogeneous perturbation effects across cell populations, DART supports efficacy-oriented hypothesis generation and highlights potential off-target effects that may be obscured in bulk analyses.

Importantly, LungChat emphasizes analytical correctness and reproducibility rather than fully autonomous hypothesis generation. Analytical steps produce provenance-tracked outputs that record the data, parameters, tools, and intermediate results required to inspect and rerun the underlying analyses. Although developed for lung biology, the architecture is tissue-agnostic, with modular agents and standardized data schemas that support extension to other organs and multi-omics domains. LungChat therefore provides an extensible system for integrated data analysis and evidence-grounded biological interpretation.

In summary, our system demonstrates how conversational multi-agent systems can serve as practical scientific interfaces that connect natural-language research questions with coordinated, reproducible computational analyses and experimentally testable hypotheses. By combining foundation models with validated analytical tools and provenance-tracked execution, LungChat provides a blueprint for transparent agentic AI systems designed to support, rather than replace, biological research.

### Use of large language models

This manuscript was written by the authors with assistance from large language models for grammar improvement, sentence clarity, and writing refinement. All scientific content, results, analyses, and conclusions were developed by the authors, who take full responsibility for the accuracy and validity of all claims.

## Acknowledgements

The authors thank the LungMAP Consortium and the contributing data-generating groups for generating and sharing the datasets analyzed in this study.

## Funding

This work was supported in part by the National Institutes of Health (NIH) National Heart, Lung, and Blood Institute (NHLBI) under Grant U24-HL148865.

## Author Contributions

P.R. led the conceptualization, design, implementation, and evaluation of the LungChat system; developed the multi-agent architecture and analytical methodology; built and integrated the frontend, backend, and tool-based analysis components; performed analyses and benchmarking; generated figures and results; and wrote the original manuscript draft. S.A. processed and analyzed spatial transcriptomics datasets, developed spatial analysis workflows and code, and contributed to spatial data interpretation. M.K. contributed to cloud infrastructure, production deployment, and operational support for the deployed system. P.L. processed and curated HLCA datasets used in the study. J.C. provided pulmonary biology expertise, scientific guidance, and feedback on the biological plausibility and interpretation of outputs. A.C. served as an AI consulting advisor and provided guidance on AI system design and implementation. V.B.S.P. contributed scientific guidance, analysis interpretation, and manuscript review and editing. B.J.A. contributed study supervision, biological interpretation, evaluation guidance, project oversight, funding acquisition, and manuscript review and editing. N.S. contributed study conceptualization, methodological guidance, evaluation design, analysis interpretation, project supervision, funding acquisition, and manuscript review and editing. All authors reviewed and approved the final manuscript.

## Data Availability

This study analyzes previously generated single-cell and spatial transcriptomics datasets available through public or consortium repositories. The Human Lung Cell Atlas v2 dataset is available through CELLxGENE under accession 6f6d381a-7701-4781-935c-db10d30de293. The fetal lung cell atlas is available through CELLxGENE under accession 2d2e2acd-dade-489f-a2da-6c11aa654028. Additional LungMAP consortium datasets are available under accessions LMEX0000004400 and LMEX0000004401. Spatial transcriptomics data are available from the Gene Expression Omnibus under accession GSE276945, including GSM8505452. Derived analysis outputs, configuration files, benchmark summaries, and figure source data generated for this study are provided in the manuscript appendices and supplementary materials, or are available upon reasonable request subject to applicable repository, consortium, and institutional restrictions.

## Code Availability

The LungChat interface described in this manuscript is deployed at https://chat.lungmap.net. The complete production application codebase, including deployment-specific frontend, infrastructure, authentication, and operational components, is not publicly released at this time. Analysis code, configuration examples, DART output tables, benchmark summaries, and other reproducibility materials sufficient to support the analyses reported in this manuscript are available upon reasonable request, subject to institutional approval.

## Ethics Statement

This study performed secondary analyses of previously generated, de-identified human single-cell and spatial transcriptomics datasets obtained from public or consortium repositories. No new human participants were recruited, and no new human specimens were collected for this study. Ethical approval and informed consent for the original data-generating studies were obtained by the respective source studies, as described in the associated publications and repository records.

## 4 Methods

### 4.1 Problem formulation and scope

We formalize the task of interactive multi-omics analysis as follows. Given a natural language query 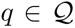, a collection of multi-omics datasets 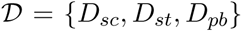 spanning single-cell RNA-seq, spatial transcriptomics, and pseudobulk expression data, and a library of analytical tools 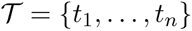, our goal is to generate a structured response 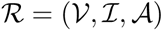 comprising visualizations 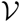, biological insights 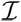, and reproducible outputs 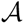.

The central challenge lies in decomposing unstructured scientific queries into executable analytical workflows while ensuring biological accuracy, cross-modal data integration, and full reproducibility. Unlike single-turn question answering, interactive analysis requires maintaining analytical state across multi-step investigations where subsequent queries build upon prior results.

### 4.2 System overview

LungChat is an AI system for integrated multi-omics analysis, designed to support biomedical discovery through interactive analysis. The system employs a hierarchical multi-agent architecture where a supervisor agent orchestrates specialized sub-agents, each augmented with domain-specific analytical tools (Figure 1). User queries are decomposed into tractable sub-tasks that execute in parallel when independent, with results synthesized into coherent scientific responses.

The architecture integrates three core capabilities: (1) multi-agent orchestration for complex query handling, (2) tool-augmented reasoning enabling precise analytical operations, and (3) unified access to heterogeneous omics data through ontology-grounded harmonization. Intermediate analytical outcomes serve as feedback signals that update the supervisor’s task decomposition, analytical resolution, and operator selection, establishing a computational lab-in-the-loop paradigm where data-driven results guide subsequent analytical actions without retraining or parameter updates.

### 4.3 Analytical workflows

#### 4.3.1 Omics datasets

LungChat operates over precompiled single-cell gene expression and spatial transcriptomics omics datasets assembled from > 37 studies curated by the LungMAP Consortium and the Human Cell Atlas initiative. Single-cell datasets are provided as CellxGene-curated `.h5ad` objects with consistent, standardized mappings to CellxGene-supported biomedical ontologies, including harmonized cell-type, disease, developmental stage, and assay annotations. To enable interactive analysis at scale, metacells and differential expression statistics are precomputed.

In addition to expression measurements, regulatory priors derived from ChIP-seq experiments and protein–protein interaction networks from prior studies are embedded locally within each `.h5ad` object using the NetPerspective software. This enables rapid retrieval of gene regulatory and interaction context during downstream analyses.

Spatial transcriptomics data include matched image-based (Xenium) and whole-transcriptome (Visium-HD) profiles. Integration between Xenium and Visium-HD datasets is performed through spot- and cell-segmented alignment and annotation stability that enables cross-platform comparison of spatial gene expression patterns and tissue organization across technologies. Appendix A provides details of the dataset curation and preprocessing undertaken.

#### 4.3.2 Specialized tools

Analytical workflows in LungChat are executed through a library of 51 specialized analytical and knowledge tools distributed across four specialist agents: the Single-Cell agent (13 tools for differential expression, visualization, and correlation analysis), Spatial agent (26 tools for spatial mapping, cell-cell communication, and neighborhood enrichment), Research agent (4 tools for functional enrichment, drug connectivity, literature search, and clinical trials), and Atlas Agent (8 tools for LungMAP/ontology access). Single-cell tools are implemented in Python using Scanpy, Matplotlib, Seaborn, and iGraph; spatial transcriptomics tools are implemented in R using Seurat, CellChat, and Squidpy. Table B1 (Appendix B) provides a complete reference for analytical, external API, and knowledge tools. At runtime these operators execute outside the language-model context as calls to dedicated containerized analysis services: single-cell tools are dispatched to a FastAPI Python sidecar (Scanpy, Matplotlib, Seaborn, iGraph) and spatial tools to an R Plumber sidecar (Seurat, CellChat, Squidpy), while user- or agent-authored custom analyses run in a separate code-execution sandbox that supports both Python and R (e.g., Seurat, tidyverse, Bioconductor) and executes as an unprivileged user. All services run as isolated Docker containers communicating over an internal bridge network authenticated by a service token, with per-request timeouts and thread/user path isolation, decoupling numerical computation from agent reasoning and from other users’ sessions.

All tools are accessed through a standardized JSON-based interface that specifies input datasets, parameter settings, and expected outputs. This abstraction enforces reproducibility and allows analytical steps to be executed deterministically. Visualization operators generate publication-ready figures in both raster (PNG) and vector (PDF) formats, accompanied by tabular outputs (TSV). For exploratory inspection of high-dimensional datasets, LungChat provides an interactive heatmap interface using Morpheus and dedicated applications for single-cell label projection (Azimuth), visualization (ShinyCell) and exploration of spatial omics (Vitessce).

#### 4.3.3 External knowledge and tools

LungChat integrates a set of API-backed external tools for functional enrichment, literature retrieval and synthesis, clinical trials, and drug connectivity analysis. Functional enrichment is performed using ToppGene, enabling pathway, disease, and drug-target annotation of gene sets. Literature reasoning is supported through the PubTator3 and EuropePMC APIs, allowing entity-aware retrieval across more than 11 million full-text PubMed Central articles and over 47 million PubMed abstracts and bioRxiv and medRxiv preprints. Clinical trial information is retrieved via the ClinicalTrials.gov API. For all external queries, input terms undergo ontology-based expansion and synonym resolution prior to submission, improving recall while preserving biological specificity. Drug Connectivity analysis is performed through iLINCS, enabling connectivity mapping between disease-associated transcriptional signatures and chemical or genetic perturbation profiles.

#### 4.3.4 Workflow control, context management, and safeguards

LungChat exposes a predefined set of workflow modes namely *Refine Question*, *Explore Data*, *Review Literature*, *Informed Analysis*, and *Test Hypothesis*. These constrain agent behavior, tool selection, and response synthesis to align with different stages of scientific inquiry. Metadata retrieval is performed in a targeted manner, loading only task-relevant fields into context to prevent overload during multi-step analytical workflows.

Short-term conversational state and intermediate workflow outputs are managed using a Redis-backed state store, enabling robust persistence across long analytical sessions. The store is a Redis Stack instance partitioned into separate logical databases for LangGraph conversation checkpoints, interface state and analytical artifacts, and per-user rate-limiting counters, with append-only persistence so that sessions survive restarts and can be resumed or replayed. To ensure analytical integrity, the supervisor agent applies layered safeguards against prompt injection and adversarial inputs, including prompt-level validation and model-based content filtering for both text and voice inputs. These controls ensure that analytical workflows remain grounded in validated tools and clearly defined scientific objectives.

### 4.4 Multi-agent architecture

#### 4.4.1 Agent roles and responsibilities

LungChat employs five specialized agents, each designed for a distinct aspect of multi-omics analysis (Table 2). The **Supervisor Agent** serves as the central orchestrator, determining optimal delegation strategies and possessing unique capabilities including dynamic analytical code execution and direct access to aggregated expression data. The **Single-Cell Agent** specializes in cell-level transcriptomic analysis, operating on pre-processed datasets with computed statistics for rapid interactive exploration. The **Spatial Agent** handles tissue-level analyses including spatial gene expression mapping and cell-cell communication inference across multiple platforms. The **Research Agent** bridges internal data with external knowledge through literature search, functional enrichment analysis, and drug-gene connectivity mapping. The **Atlas Agent** provides structured LungMAP/ontology access to domain-specific resources and controlled vocabularies.

**Table 2:** Agent specifications.

| Agent | Role | Analytical Scope |
| --- | --- | --- |
| Supervisor | Query routing, result synthesis, code execution | Cross-modal coordination |
| Single-Cell | Differential expression, cell characterization | scRNA-seq transcriptomics |
| Spatial | Tissue organization, cell communication | Spatial transcriptomics |
| Research | Literature integration, enrichment, connectivity | External knowledge |
| Atlas | Ontology queries, vocabulary lookup | Domain databases |

#### 4.4.2 Inter-agent communication and execution model

Agent coordination follows a stateless delegation model designed for reproducibility and modularity. Subagents execute independently without access to conversation history or other agents’ intermediate reasoning; they receive only the delegated task specification and return self-contained responses. This design limits the influence of unrelated conversation history and improves the consistency and reproducibility of subagent execution.

The supervisor implements parallel task execution for independent analytical operations. When a query decomposes into non-dependent sub-tasks, these execute concurrently, significantly reducing response latency. Dependent tasks execute sequentially with results from prior steps informing subsequent operations. Task execution follows a managed lifecycle with explicit states: *pending*, *in_progress*, and *completed*. Failed tasks trigger supervisor-mediated retry with parameter adjustment before graceful degradation with informative error reporting.

To support inter-agent collaboration and long-horizon workflows, LungChat employs a shared virtual filesystem serving as a common workspace for intermediate results and analysis outputs. When tool executions produce outputs exceeding a configurable size threshold, results are automatically materialized as filesystem outputs and replaced in context with lightweight file references. All agents have read and write access to the shared workspace, enabling downstream agents to inspect, extend, or refine outputs produced by upstream analyses. The workspace is thread-scoped: each session writes to an isolated output namespace, and file access is mediated by a filesystem backend that resolves artifact references, blocks path traversal and symbolic-link escapes, and enforces size limits, preserving per-user and per-thread isolation across concurrent sessions.

The stateless delegation model serves a dual purpose beyond modularity: it implements context engineering principles that mitigate known limitations of long-context language model reasoning [15]. Each specialist agent operates in a scoped context window containing only its domain-relevant tools and instructions, preventing the cross-domain parameter leakage and instruction dilution that arise when a single agent must reason over all 51 tools simultaneously. Sub-agents are invoked without conversation history, ensuring that reasoning quality at turn *n* of a session is structurally independent of prior turns, a property we term *stale-context resistance*. This design aligns with recent findings that multi-agent collaboration through scoped worker contexts outperforms monolithic single-agent approaches on long-context tasks [30] and recommended practices for managing context degradation in agentic systems [2].

### 4.5 Tool-augmented reasoning

#### 4.5.1 Tool abstraction layer

LungChat augments language model reasoning with a structured library of analytical tools organized into four functional operator classes: **Analysis Operators** perform statistical and computational procedures including ToppGene for functional enrichment testing and iLINCS for drug-gene perturbation connectivity analysis, alongside differential expression and pathway overlap operators; **Visualization Operators** generate publication-ready figures in both raster and vector formats; **Knowledge Operators** interface with external resources including biomedical literature databases, clinical trial registries, and gene annotation services; **Data Access Operators** provide structured access to consortium data APIs and curated omics datasets.

#### 4.5.2 Hybrid tool selection mechanism

With 51 tools across 4 operator classes, exhaustive evaluation of all tools for each query is computationally prohibitive and degrades response quality through irrelevant context. We implement a two-stage hybrid selection mechanism balancing broad recall with precise final selection.

Given a query *q* and tool library 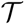, the selection process is:

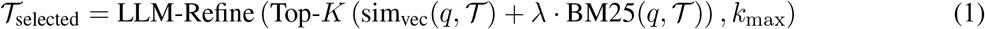

where sim_vec_ computes semantic embedding similarity and BM25 provides lexical keyword matching. In practice, *λ* = 0.3 balances semantic and lexical signals, and *K* = 15 defines the initial candidate pool. The candidate pool is then refined by a lightweight language model to select the final *k*_max_ tools (typically 3) based on query-tool relevance reasoning. This hybrid approach captures both semantic similarity for paraphrased queries and lexical overlap for specific technical terms, substantially reducing token consumption while maintaining selection accuracy.

### 4.6 Multi-omics data integration

#### 4.6.1 Data modalities

LungChat integrates three complementary omics modalities through a unified query interface, as summarized in Table 1. **Single-cell RNA-seq** data comprises 39 datasets totaling 2.8 million cells spanning fetal development, healthy adult tissue, and 15 disease contexts including idiopathic pulmonary fibrosis, COPD, and COVID-19, with pre-computed differential expression statistics and harmonized cell type annotations across common expression schemas. **Spatial transcriptomics** data includes IPF dataset across both Visium-HD and Xenium platform types capturing spatially-resolved gene expression with pre-computed cell-cell communication objects. **Pseudobulk expression** data provides aggregated views across cell types and conditions in a queryable warehouse with comprehensive differential expression signatures supporting cross-dataset comparisons.

#### 4.6.2 Ontology-grounded metadata harmonization

Metadata harmonization is performed by mapping heterogeneous author-provided annotations to standardized biomedical ontologies including CL, UBERON, MONDO, EFO, and HANCESTRO. This approach achieves robust resolution of heterogeneous annotations to canonical ontology terms, enabling consistent cross-dataset queries. All ontology mappings are versioned and documented to ensure consistent interpretation. Author-provided labels are resolved to canonical ontology identifiers and stored alongside the harmonized expression warehouse in a SQLite catalog with full-text (FTS5) indices over cell-type, disease, assay, developmental-stage, sex, and ancestry vocabularies, enabling fuzzy and synonym-aware term matching. At query time, metadata fields are retrieved on demand and search terms are expanded across ontology ancestors and descendants before warehouse or external-API submission, so that a single disease or cell-type term transparently captures its subtypes and related annotations across heterogeneously labeled datasets.

### 4.7 Direction-Aware Repurposing and Targeting (DART)

To enable cell-type-resolved therapeutic prioritization, we introduce DART, a method that distinguishes perturbations reversing disease-associated transcriptional programs from those that reinforce them. Unlike conventional connectivity analyses producing compound-level scores aggregated across heterogeneous cell populations, DART preserves cell-type resolution throughout analysis, allowing antagonistic effects across lineages to be explicitly identified.

#### Directional scoring

Given a disease differential expression signature 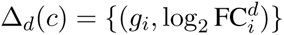 for cell type *c* and a perturbation-induced transcriptional signature 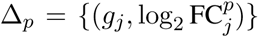, DART converts fold changes to binary directional indicators (+1 for upregulation, −1 for downregulation). For overlapping genes *G*_overlap_ = *G_d_* ∩ *G_p_*, genes are classified by directional concordance:

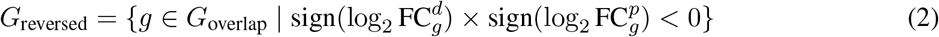

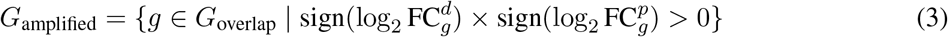

DART computes two heuristic metrics: Pearson correlation on binary directions *ρ*(*c*) = corr_Pearson_(dir*_d_,* dir*_p_*) and reversal rate *R*(*c*) = |*G*_reversed_|*/*|*G*_overlap_|, where negative *ρ* and high *R* indicate therapeutic reversal. Perturbations are ranked by reversal rate (reversal mode) or its complement (amplification mode), enabling prioritization of compounds that oppose or reinforce disease programs. Directional stratification by disease-upregulated versus disease-downregulated genes enables cell-type-specific safety profiling. DART is implemented as a composable analysis operator operating on precomputed signatures and can be applied wherever matched disease and perturbation transcriptional profiles are available.

##### Algorithm 1

**Query Decomposition and Execution**

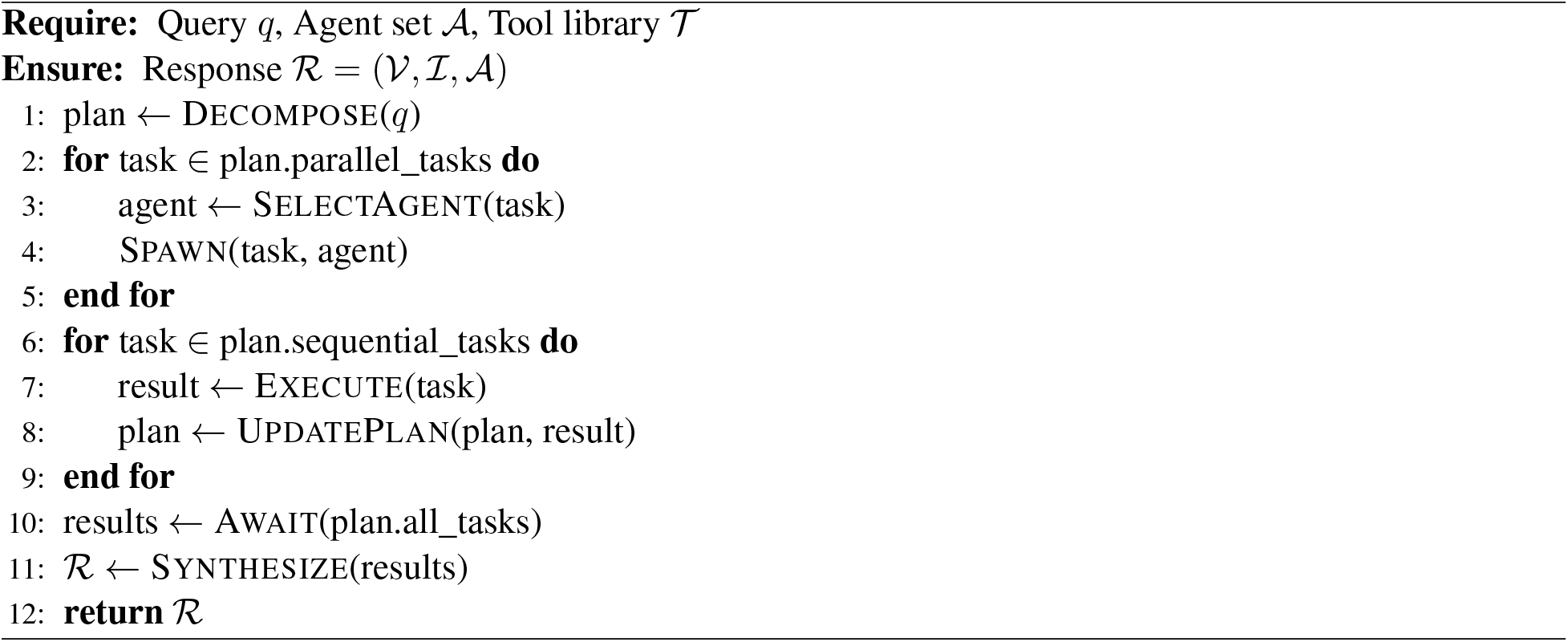

### 4.8 Iterative scientific workflows

#### 4.8.1 Planning and task decomposition

Complex scientific queries require structured decomposition into executable sub-tasks. LungChat implements explicit planning through a task decomposition mechanism that transforms natural language queries into coordinated analytical workflows, see Algorithm 1.

The Decompose function employs few-shot prompting with examples of query-to-task mappings. SelectA-gent uses rule-based routing based on task keywords with LLM fallback for ambiguous cases. The planning mechanism identifies task dependencies to maximize parallel execution. The supervisor can dynamically revise plans based on intermediate results - if initial analysis reveals unexpected patterns, subsequent tasks can be adjusted or additional analyses spawned without requiring user intervention.

#### 4.8.2 Planning tool

Complex biomedical workflows often require significant computational time, creating a "black box" latency period that erodes user trust. To address this, LungChat implements a real-time task synchronization layer. When executing multi-step protocols, the Supervisor agent actively maintains a structured state object (todos) that maps the analysis plan to visible UI elements. This state is synchronized via a low-latency Redis layer to the frontend, transforming the waiting period into an interactive progress log. Users see the agent decompose their query into discrete biological tasks (e.g., "1. Query Omics Warehouse", "2. Perform Enrichment"), mark them as `in_progress`, and resolve them to completed in real-time. This mechanism provides immediate visual verification that the agent is executing the intended logic, distinguishing productive "reasoning time" from system stalls.

#### 4.8.3 Provenance-tracked reproducibility and memory

Scientific analysis requires reproducibility beyond context. LungChat structures all outputs as persistent outputs with complete provenance: **Configuration Outputs** capture all input parameters, data filters, and analysis settings as JSON; **Visualization Outputs** include figures in multiple formats (PNG, PDF) with accompanying data tables (TSV); **Execution Traces** document the sequence of analytical operations, agent delegations, and intermediate results; **Session Reports** provide automated summaries of analytical work-flows and key findings, see Appendix C for an example. In addition, each analytical step emits a UID (code_used__*_.py or code_used__*_.R) containing the exact implementation source and the resolved parameters used for that run, so that any figure or table can be regenerated independently of the conversational interface. Code, configurations, and input files can be downloaded and re-run by the user locally to reproduce results or the JSON directly supplied back to LungChat to replicate or modify prior analyses.

Beyond single-session analysis, LungChat maintains user-specific long-term analytical context across sessions, storing structured research state including previously analyzed genes, selected cell types, and references to generated outputs. Memory retrieval is scoped to analytical context and does not influence model parameters or tool execution logic, ensuring reproducibility while enabling cumulative scientific workflows. Extended analyses exceeding context capacity employ dynamic compression using a dedicated summarization module, preserving recent interactions while compressing prior context. Combined with filesystem-based eviction of large tool outputs, this strategy decouples analytical data volume from context length.

### 4.9 Implementation and inference regime

LungChat operates entirely at inference time using frozen foundation models. No task-specific training, fine-tuning, or gradient updates are performed on any model components. All capabilities including multi-agent coordination, tool selection, and response synthesis emerge from prompt engineering, structured tool augmentation, and orchestration logic. The system employs Claude Sonnet 4.6 and GPT-4 class models with long-context windows (≥128,000 tokens). Tool selection uses low-temperature decoding (set 0.1), while response synthesis uses moderate temperature (set 0.7). This design enables rapid capability updates through prompt and tool modifications without retraining, while leveraging the broad knowledge encoded in pre-trained foundation models.

## A Dataset curation and preprocessing

### A.1 Single-cell RNA-seq

Four independent single-cell RNA-Seq collections were integrated into LungChat’s corpus of initial available datasets. The largest of these dataset collections consist of 2.2 million cells from 486 lungs from Human Lung Cell Atlas (HLCA) version 2 [25] object in CellxGene (accession: 6f6d381a-7701-4781-935c-db10d30de293 https://cellxgene.cziscience.com/collections/6f6d381a-7701-4781-935c-db10d30de293). HLCA is a collection of reprocessed raw droplet sequencing data 35 single-cell studies, spanning 14 distinct disease entities. To efficiently assess and analyze this data, the R library SuperCell v.1.0 was used to produce 50,520 metacells from the HLCA, separately for each individual donor and HLCA consortia annotated cluster (ann_finest_level). A lung cell atlas of fetal development was also obtained from CellxGene (accession: 2d2e2acd-dade-489f-a2da-6c11aa654028 https://cellxgene.cziscience.com/collections/2d2e2acd-dade-489f-a2da-6c11aa654028), spanning 5-22 post-conception weeks [8]. Two additional LungMAP consortia datasets were provided to LungChat, comprising infant lungs at the time of death from bronchopulmonary dysplasia or other causes (controls) (LungMAP.net accession: LMEX0000004400 https://www.lungmap.net/dataset/?dataset_id=LMEX0000004400, LMEX0000004401 https://www.lungmap.net/dataset/?dataset_id=LMEX0000004401) [24]. Cell annotations for each dataset include author curated at different levels of resolution and Cell Ontology unique IDs. All datasets are formatted by LungChat pre-processing as anndata H5AD objects, from source H5AD or RDS, using developed automated processing workflows (GitHub). This workflow retains author provided UMAP coordinates (averaged for metacells) and harmonized sample metadata (CellxGene). The preprocessing workflow further produces normalized expression using the Scanpy Python package (natural log of counts per 10,000 reads), translates Ensembl primary identifiers to gene symbols (Ensembl 100), computes cell population marker gene and covariate differential gene expression versus appropriate control conditions (wilcoxon, FDR corrected) and retains DEGs with fold *>* 1.2 and Mann-Whitney U test *p <* 0.05 (FDR adjusted) in uns["rank_genes_groups"]. Study-specific gene signatures are stored for later comparison in LungChat. Each anndata object is augmented with a database of protein-protein and protein-DNA interactions from the AltAnalyze NetPerspective database [28], for fast retrieval and display of inferred gene regulatory networks in LungChat.

### A.2 Spatial transcriptomics

Image-based (10x Genomics Xenium) and whole transcriptome (10x Genomics Visium-HD) spatial transcriptomics from a collection of human healthy and IPF lungs were obtained from the Gene Expression Omnibus database (GSE276945 https://www.ncbi.nlm.nih.gov/geo/query/acc.cgi?acc=GSE276945). Each patient sample corresponds to a region of interest (ROI), tiled on the Xenium (343 gene probes) or Visium slide (18,085 genes, GSM8505452 https://www.ncbi.nlm.nih.gov/geo/query/acc.cgi?acc=GSM8505452). For Xenium samples (n=35), author generated cell segmentation (1.6 million cells), curated cell annotation (Final_CT) and niche annotation (TNiche and CNiche) were retained, along with sample level metadata (i.e., sex, age). Segmented cell annotations were augmented with Robust Cell Type Decomposition (RCTD) from the spacexr package that was applied to derive supervised cell annotations for RCTD-classified singlets using the human CellRef transcriptome as a reference. For Visium-HD, as no author annotation for 8*µ*m spots were provided, clusters, the same RCTD protocol was applied for Visium spots, to generate annotations.

To derive hybrid multi-modal cell annotations from the integration of matching Xenium and Visium samples, we reoriented, scaled and aligned Xenium segmented map (query) to Visium (reference) coordinates, considering unique cell/spot correspondences (dplyr package in R). For downstream integrated analyses, all cell/spots were downsampled to those with 1:1 correspondences using the find nearest neighbor function (FNN) package and get.knnx() function. Mapping was verified using COL1A1 expression, which was highly expressed by both platforms. Cluster annotations were mapped across 1:1 mapped cells/spots across platforms to define the most stable cell populations using a new spatial library within the scTriangulate package (https://github.com/frankligy/scTriangulate). scTriangulate [14] uses a game-theory approach to compare distinct stability metrics (marker gene expression, reclassification) for each single-cell cluster, separately for each modality evaluated. In this instance, all Xenium and Visium cell or niche clusters were evaluated using the integrated object, with RNA values and spatial coordinates from each platform considered as separate modalities. A final set of 29 multimodal stable clusters were defined from this analysis and stored in the integrated RDS object for LungChat analysis.

To infer cell-cell communication networks, we applied the python package Squidpy to assess cell-type spatial proximity via neighborhood enrichment (z-score) and CellChat to infer cell-cell and receptor-ligand interactions. These analyses were pre-computed using the Xenium dataset considering control (less affected) and IPF (more affected) samples. Both Squidpy and CellChat were applied to different annotation layers, namely author (Final_CT), cellular niches (CNiche, TNiche) and Final_CT annotations that subdivide individual niches. Computed results were stored in Seurat RData objects for direct access by LungChat.

### B Tool reference and output specifications

### B.1 Tool scope

This appendix provides a capability-level reference of analytical, external API, and knowledge tools used by the Supervisor, Single-Cell, Spatial, and Research agents (Atlas Agent excluded).

### B.2 Capability-level tool catalog

**Table B1:** Capability-level reference of analytical, external API, and knowledge tools used by the Supervisor, Single-Cell, Spatial, and Research agents (Atlas Agent excluded), including output formats and descriptions.

| Tool | Output format | Description |
| --- | --- | --- |
| **Supervisor Agent Tools** |  |  |
| query_pseudobulk_expression_data_tool | JSON | Query pseudobulk omics warehouse (DEG lookup, aggregation, comparisons, signature overlap, drug-cell-type impact) |
| execute_code_tool | JSON | Run Python or R in sandbox |
| **Single-Cell (scRNA-seq) Agent Tools** |  |  |
| plot_heatmap | JSON, PDF, PNG, TSV | Clustered heatmap of gene expression across cell types/conditions. |
| plot_radar | JSON, PDF, PNG | Radar/spider chart of cell type composition across conditions. |
| plot_cell_frequency | JSON, PDF, PNG | Box/violin of cell frequencies per donor; Mann-Whitney U. |
| plot_volcano | JSON, PDF, PNG | Volcano: log2FC vs FDR; top genes labeled. |
| plot_dotplot | JSON, PDF, PNG | Dot plot: % expressing (size), mean expression (color) across cell types. |
| plot_violin | JSON, PDF, PNG | Violin of one gene expression across cell types/conditions. |
| plot_venn | JSON, PDF, PNG | Venn diagram of gene set overlaps (2–3 sets). |
| plot_upset_genes | JSON, PDF, PNG | UpSet plot for 4+ gene set overlaps. |
| plot_umap | JSON, PDF, PNG | UMAP 2D embedding; optional gene overlay. |
| plot_network | JSON, PDF, PNG | Gene–gene interaction/regulatory network. |
| plot_gene_correlation | JSON, PDF, PNG, TSV | Pearson correlation of genes with query gene. |
| generate_stats | JSON, TSV | Export DEG/marker statistics. |
| **Spatial Agent Tools** |  |  |
| plot_spatial | JSON, PDF, PNG | Spatial plot of cell annotations (Final_CT, TNiche, CNiche, etc.) on coordinates. |
| plot_visualize_celltypes_spatial_distribution | JSON, PDF, PNG | Multi-panel spatial distribution of cell types across samples/groups. |
| plot _ compare _ spatial _ rna _ with _ histology _ quadpanel | JSON, PDF, PNG | 4-panel: Visium-HD, Xenium, H&E, DAPI. |
| plot_umap | JSON, PDF, PNG | UMAP colored by annotation (spatial data). |
| plot_visualize_gene_expression_spatial | JSON, PDF, PNG | Gene expression on spatial coordinates (Xenium/Visium). |
| plot_barchart_top_correlated_visium_xenium_rna | JSON, PDF, PNG | Bar chart of top correlated genes across Visium/Xenium. |
| plot_evaluate_cluster_stability_report | JSON, PDF, PNG | Report on annotation stability across sources. |
| rank_celltype_marker_genes_by_specificity | JSON, TSV | Rank cell types by marker specificity (t-test/FDR). |
| plot_celltype_marker_volcano | JSON, PDF, PNG | Volcano of marker genes for one cell type. |
| plot_compare_markers_across_platforms_dotplot | JSON, PDF, PNG | Dot plot comparing markers across platforms. |
| plot_evaluate_annotation_concordance_dotplot | JSON, PDF, PNG | Dot plot of annotation concordance. |
| plot_scatter_matched_xenium_visium_spots | JSON, PDF, PNG | Scatter of matched Xenium-Visium spot expression. |
| plot_differential_expression_volcano | JSON, PDF, PNG | Volcano for spatial DEGs between groups. |
| plot_cellchat_chord_diagram | JSON, PDF, PNG | CellChat chord diagram of signaling. |
| plot_cellchat_heatmap | JSON, PDF, PNG | CellChat signaling strength heatmap. |
| plot_cellchat_signaling_pathway_network | JSON, PDF, PNG | CellChat pathway network. |
| extract_cellchat_interactions_subset | JSON, TSV | Extract subset of CellChat interactions. |
| plot_squidpy_neighbour_enrichment_heatmap | JSON, PDF, PNG | Squidpy neighborhood enrichment heatmap. |
| plot_squidpy_celltype_neighborhood_enrichment | JSON, PDF, PNG | Squidpy cell type neighborhood enrichment. |
| plot_identify_significant_proportion_changes | JSON, PDF, PNG, TSV | Proportion changes between groups. |
| plot_compare_different_cell_annotations_heatmap | JSON, PDF, PNG | Heatmap comparing two annotation columns. |
| plot_identify_celltypes_enriched_in_niches | JSON, PDF, PNG | Cell types enriched in niches. |
| plot_squidpy_compare_neighborhood_changes | JSON, PDF, PNG | Compare Squidpy neighborhoods across groups. |
| identify_unique_cellchat_interactions_spatial | JSON, TSV | Unique CellChat interactions per group. |
| plot_compare_communication_profiles | JSON, PDF, PNG | Compare CellChat communication between groups. |
| **Research Agent Tools** |  |  |
| gene_functional_enrichment_tool | JSON, TSV | ToppGene enrichment (20 categories). |
| literature_search_tool | JSON | PubMed + Europe PMC; ranking, preprints, entity extraction. Returns papers, insights. |
| clinical_trials_search_tool | JSON | ClinicalTrials.gov v2; term expansion (MONDO/EFO). Search or retrieve by NCT ID. |
| drug_connectivity_tool | JSON, PDF, PNG, TSV | iLINCS: signature connectivity, compound investigation, gene knockout, similar compounds. |
| web_search_tool | JSON | Google Serper API; fallback for non-biomedical queries. Returns title, snippet, link. |
| think_tool | JSON | Record strategic reflection (progress, gaps, next steps). Returns confirmation. |

## C Provenance-tracked configuration example

**Figure C1:**
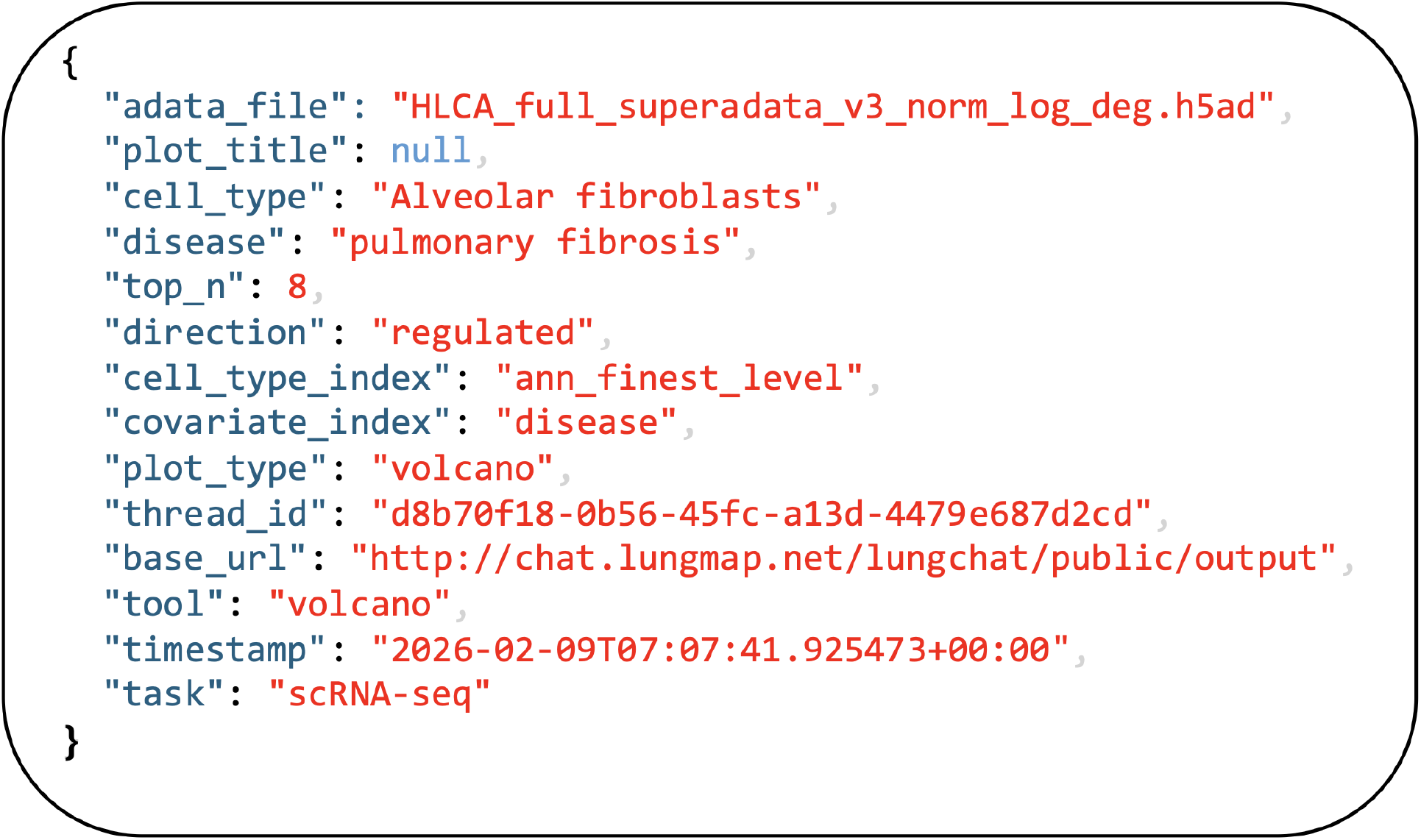
Provenance-tracked configuration output for reproducible tool execution. Example JSON payload automatically saved by LungChat for a differential-expression volcano plot request (dataset, cell type, disease, direction, and tool parameters), enabling deterministic re-execution and auditability of analysis settings as described in Section 4.8.3.

## D Capability comparison with related systems

**Table D1:**
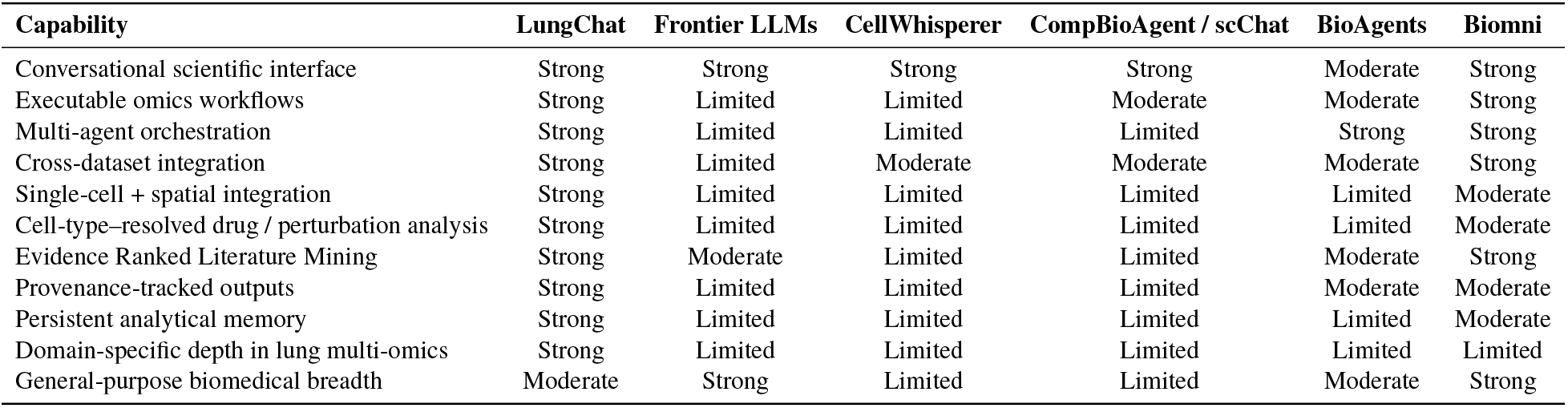
Capability-level comparison with related conversational and agentic biomedical AI systems.

| Capability | LungChat | Frontier LLMs | CellWhisperer | CompBioAgent / scChat | BioAgents | Biomni |
| --- | --- | --- | --- | --- | --- | --- |
| Conversational scientific interface | Strong | Strong | Strong | Strong | Moderate | Strong |
| Executable omics workflows | Strong | Limited | Limited | Moderate | Moderate | Strong |
| Multi-agent orchestration | Strong | Limited | Limited | Limited | Strong | Strong |
| Cross-dataset integration | Strong | Limited | Moderate | Moderate | Moderate | Strong |
| Single-cell + spatial integration | Strong | Limited | Limited | Limited | Limited | Moderate |
| Cell-type-resolved drug / perturbation analysis | Strong | Limited | Limited | Limited | Limited | Moderate |
| Evidence Ranked Literature Mining | Strong | Moderate | Limited | Limited | Moderate | Strong |
| Provenance-tracked outputs | Strong | Limited | Limited | Limited | Moderate | Moderate |
| Persistent analytical memory | Strong | Limited | Limited | Limited | Limited | Moderate |
| Domain-specific depth in lung multi-omics | Strong | Limited | Limited | Limited | Limited | Limited |
| General-purpose biomedical breadth | Moderate | Strong | Limited | Limited | Moderate | Strong |

Direct quantitative benchmarking against competing systems is precluded by non-overlapping analytical capabilities: the end-to-end workflows evaluated here, spanning differential expression, functional enrichment, drug connectivity screening, cell-type–resolved perturbation analysis (DART), and spatial transcriptomics, require integrated tool access that no single competing system currently supports. While several systems offer public deployments (Biomni, CellWhisperer, CompBioAgent), each addresses a subset of these capabilities (Table D1). We therefore provide this qualitative capability comparison alongside a controlled single-agent vs. multi-agent architecture ablation (Section 2.2).

## E DART case studies

This appendix provides comprehensive execution provenance for the two DART case studies described in Section 2.3: (1) the IPF Saracatinib discovery, executed as a single-prompt, zero-shot workflow; and (2) the COPD Fluticasone Propionate discovery, executed as a single-prompt, cell-type-resolved zero-shot workflow with per-cell-type iLINCS screening. Both case studies were conducted in fresh sessions with no prior conversation history. All data, tables, and narrative text below are derived directly from the LungChat conversation exports and LangSmith agent execution traces. Figures reference generated outputs archived with the session.

### E.1 IPF: Saracatinib Discovery

#### Supplementary E1. IPF DART Case Study: Single-Prompt Zero-Shot Therapeutic Discovery

##### User prompt (verbatim)

Identify therapeutic targets and drug repurposing candidates for IPF based on cell-type–specific disease programs in fibroblast populations most associated with fibrosis, comparing diseased and healthy states within those cells, and how these manifest across lung cell types and tissue context, including safety. Prioritize drugs that reverse this program, and for the top candidate, assess its cell-type-specific effects across the lung.

##### Supervisor decomposition

LungChat’s supervisor agent decomposed this single natural-language prompt into six coordinated sub-tasks, executed across three specialized agents with parallel dispatch for independent operations:

**1. Define the disease program:** DEGs in IPF fibroblasts vs. healthy controls (HLCA)
**2. Drug repurposing:** iLINCS screen for compounds reversing the fibroblast signature
**3. Pathway enrichment:** Characterize the fibrotic program biologically (ToppGene)
**4. Cell-type impact:** Map the top drug candidate across all lung cell types (DART)
**5. Spatial context:** Fibroblast niches and ligand-receptor signaling in IPF tissue
**6. Safety synthesis:** Integrate all findings into a prioritized report

**Steps 1, 3, and 5 were dispatched in parallel. Steps 2 and 4 were executed sequentially after Step 1 completed.**

##### Complete execution trace

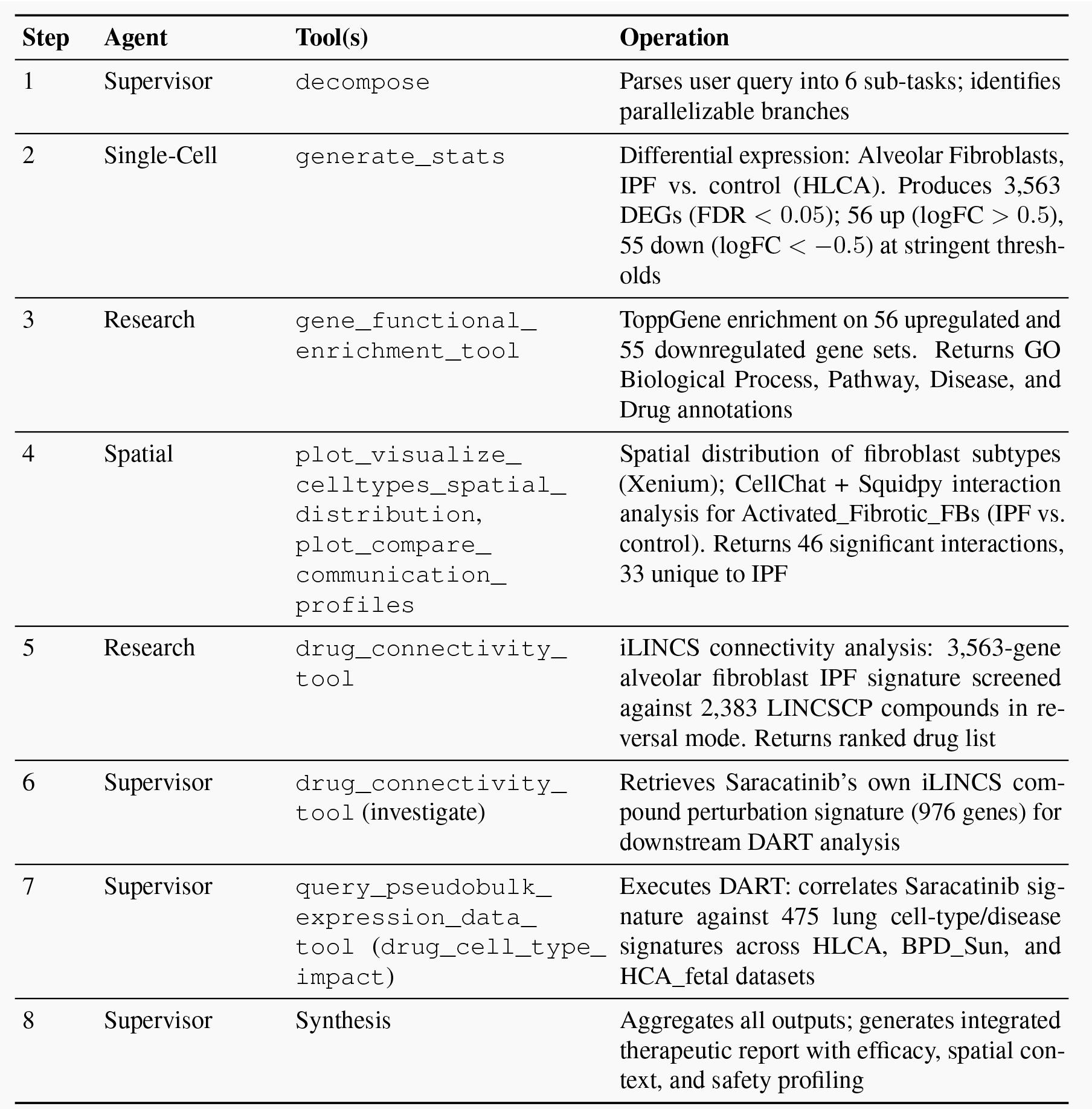

##### Step 2 output: fibroblast disease program

Analysis of alveolar fibroblasts from the HLCA identified a robust fibrotic gene expression signature comprising **3,563 significant DEGs** (FDR *<* 0.05). The upregulated program is dominated by extracellular matrix remodeling, cell adhesion, and pro-inflammatory signaling; the downregulated program indicates loss of fibroblast detoxification and quiescence capacity.

| Gene | Log2FC | Functional Role |
| --- | --- | --- |
| IL13RA2 | +9.69 | IL-13 decoy receptor; fibrosis amplifier |
| THY1 | +6.54 | Fibroblast activation marker (CD90) |
| NFATC2 | +5.63 | Transcription factor; myofibroblast activation |
| CPXM1 | +5.59 | Extracellular metalloprotease |
| IGFBP4 | +5.37 | IGF signaling modulator |
| SPHK1 | +4.87 | Sphingosine kinase; pro-fibrotic lipid signaling |
| IL32 | +4.87 | Pro-inflammatory cytokine |
| SAMD11 | +4.84 | Transcriptional regulator |
| SERPINE2 | +4.79 | Serine protease inhibitor; ECM remodeling |
| RUNX1 | +3.79 | Myeloid/fibroblast TF; differentiation driver |

*Top Downregulated Genes (Lost Homeostatic Program):*
| Gene | Log2FC | Functional Role |
| --- | --- | --- |
| CYP4B1 | −32.6 | Xenobiotic metabolism; lung-specific detoxification |
| SAA1 | −6.12 | Acute phase protein |
| NKD1 | −5.94 | Wnt pathway negative regulator |
| HIF3A | −5.40 | Hypoxia response modulator |
| CCN5 | −3.55 | Anti-fibrotic matricellular protein |
| ALDH1A1 | −3.14 | Retinoid metabolism; fibroblast quiescence |
| PDGFRA | −1.59 | PDGF receptor; fibroblast identity marker |

##### Step 3 output: functional enrichment

ToppGene enrichment of the 56 upregulated genes identified the following top-ranked processes and pathways:

| Term | p-value | Genes |
| --- | --- | --- |
| Regulation of cell adhesion | $2.47 \times 10^{-9}$ | 15/49 |
| Regulation of cell migration | $1.36 \times 10^{-8}$ | 16/49 |
| Regulation of cell motility | $2.98 \times 10^{-8}$ | 16/49 |
| Positive regulation of cell adhesion | $1.06 \times 10^{-7}$ | — |
| Regulation of plasminogen activation | $1.08 \times 10^{-7}$ | — |
| Dissolution of fibrin clot (Reactome) | $7.69 \times 10^{-6}$ | — |
| Eicosanoid synthesis (WikiPathways) | $4.06 \times 10^{-5}$ | — |
| HIF-1 transcription factor pathway | $4.96 \times 10^{-5}$ | — |
| RUNX1-regulated myeloid differentiation | $1.94 \times 10^{-4}$ | — |

The downregulated program was enriched for Phase I xenobiotic metabolism (CYP4B1, FMO2, INMT; *p* = 6.01 × 10^−8^), amino acid metabolism, and RECK pathway (MMP regulation), indicating loss of normal fibroblast detoxification capacity.

**Key druggable targets** identified by enrichment: kinases (MAPKAPK2, PIM3, NEK6), receptors (IL1R1, TNFRSF1A, LPAR1, AGTR1), and transcription factors (RUNX1, NFATC1/NFATC2, SOX4, FOXP1).

**Figure E1:**
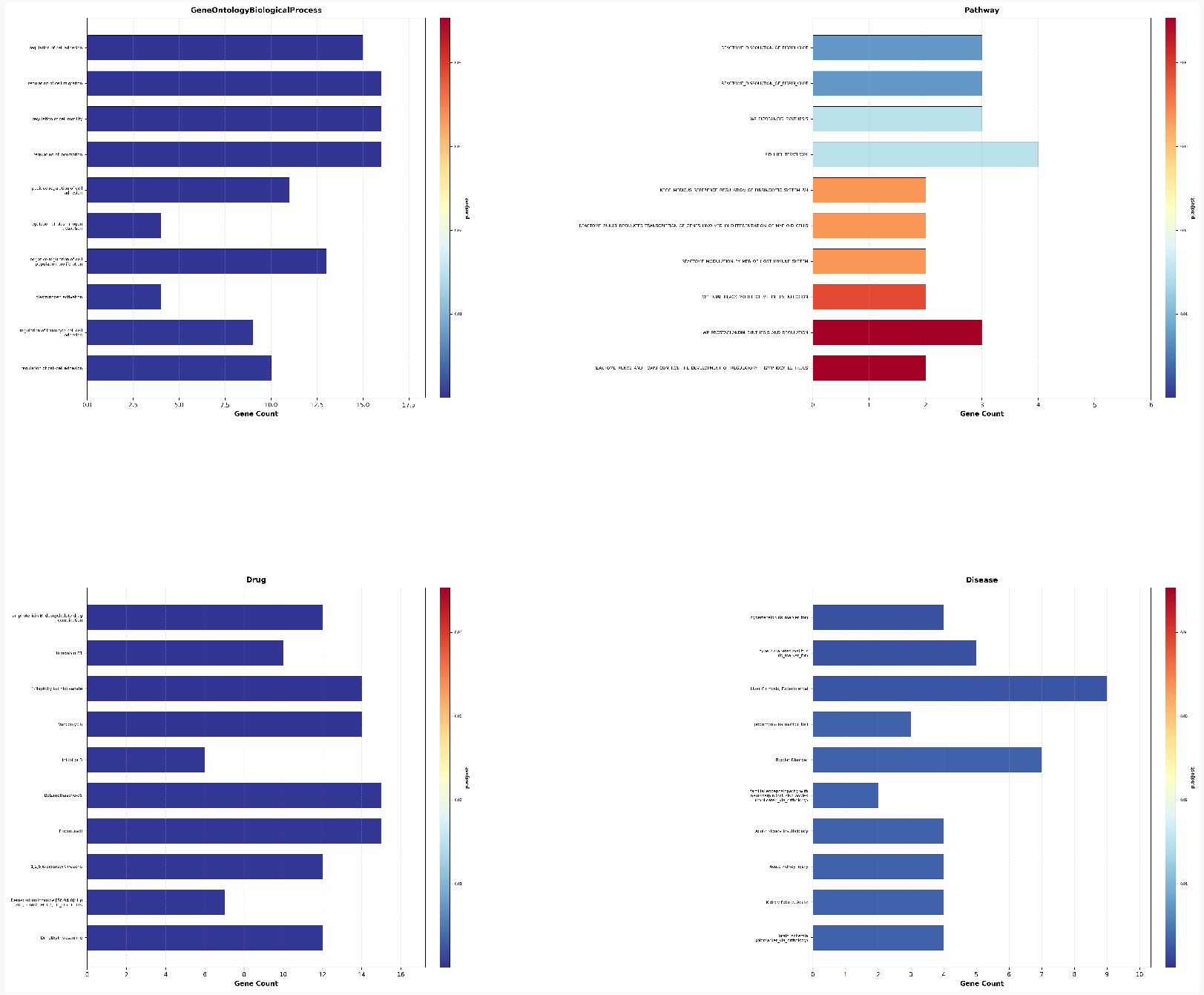
ToppGene functional enrichment of the top 56 upregulated IPF alveolar fibroblast genes. GO Biological Process terms highlight cell adhesion, migration, and HIF-1 signaling; Disease ontology maps to pulmonary fibrosis and related pathologies; Drug associations support kinase inhibitor candidates. All terms FDR-corrected, *q <* 0.05.

##### Step 4 output: spatial context

Spatial transcriptomics (Xenium, single-cell resolution) revealed seven distinct fibroblast subtypes with segregated niches in unaffected lung tissue: Alveolar_FBs distributed throughout parenchyma, Subpleural_FBs at tissue edges, Myofibroblasts in focal clusters, and Activated_Fibrotic_FBs near remodeling zones. CellChat and Squidpy interaction analysis of Activated_Fibrotic_FBs (IPF vs. control) identified:

- 4**6 significant interactions** in More_Affected tissue; **33 unique to IPF**
- **Gained in IPF:** Interactions with KRT5^−^/KRT17^+^ aberrant basaloid cells and capillary populations; monocyte–fibroblast crosstalk amplified
- **Lost in IPF:** Lymphatic and venous endothelial communication; NK cell interactions reduced
- **Top ligand–receptor pairs:** VEGFA–VEGFR2, EREG–EGFR, SPP1–ITGAV/ITGB1, SPP1–CD44, CCL2–ACKR1

**Figure E2:**
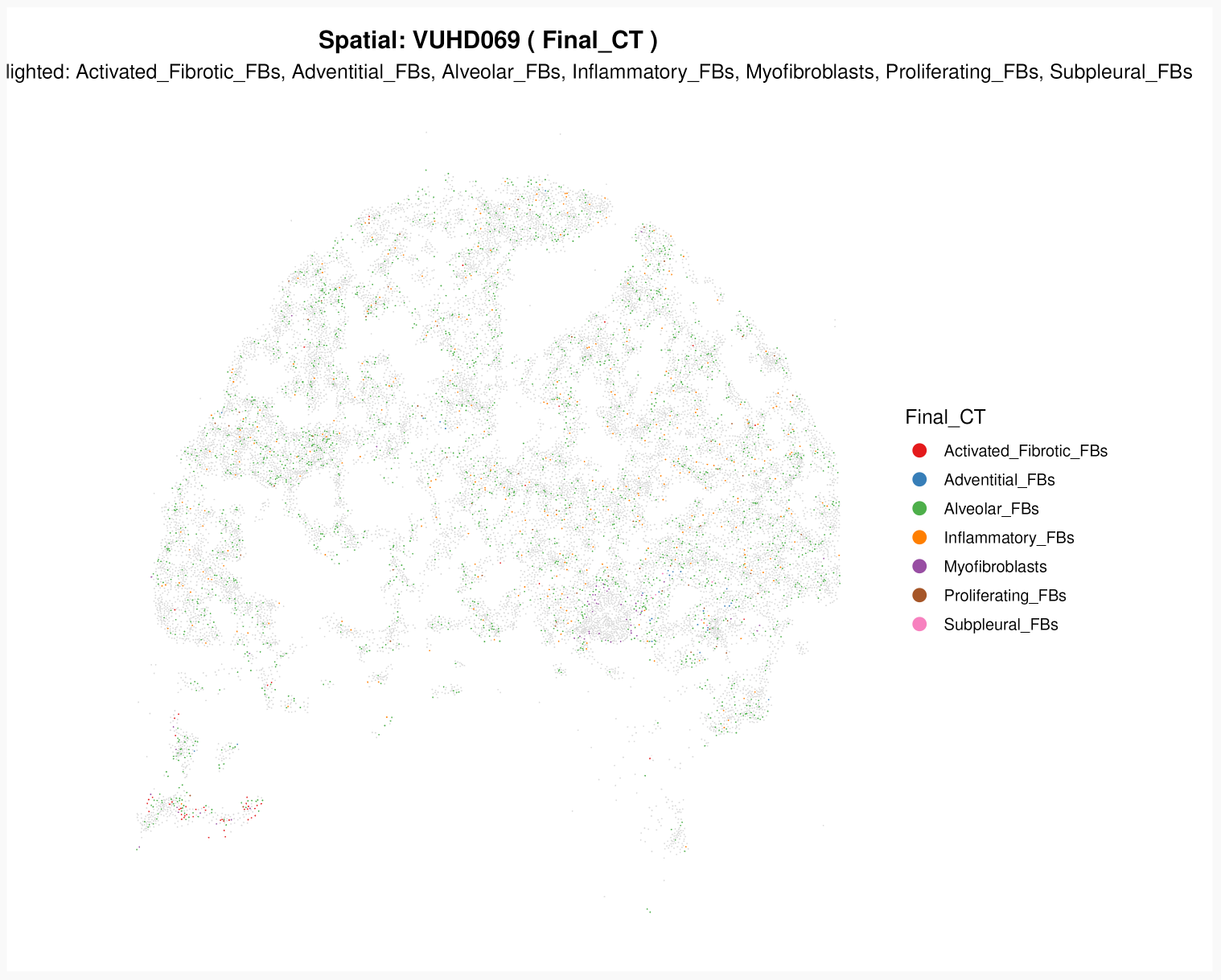
Spatial cell-type distribution in Unaffected IPF tissue (Xenium). UMAP overlay reveals fibroblast subpopulation geography across seven distinct niches.

**Figure E3:**
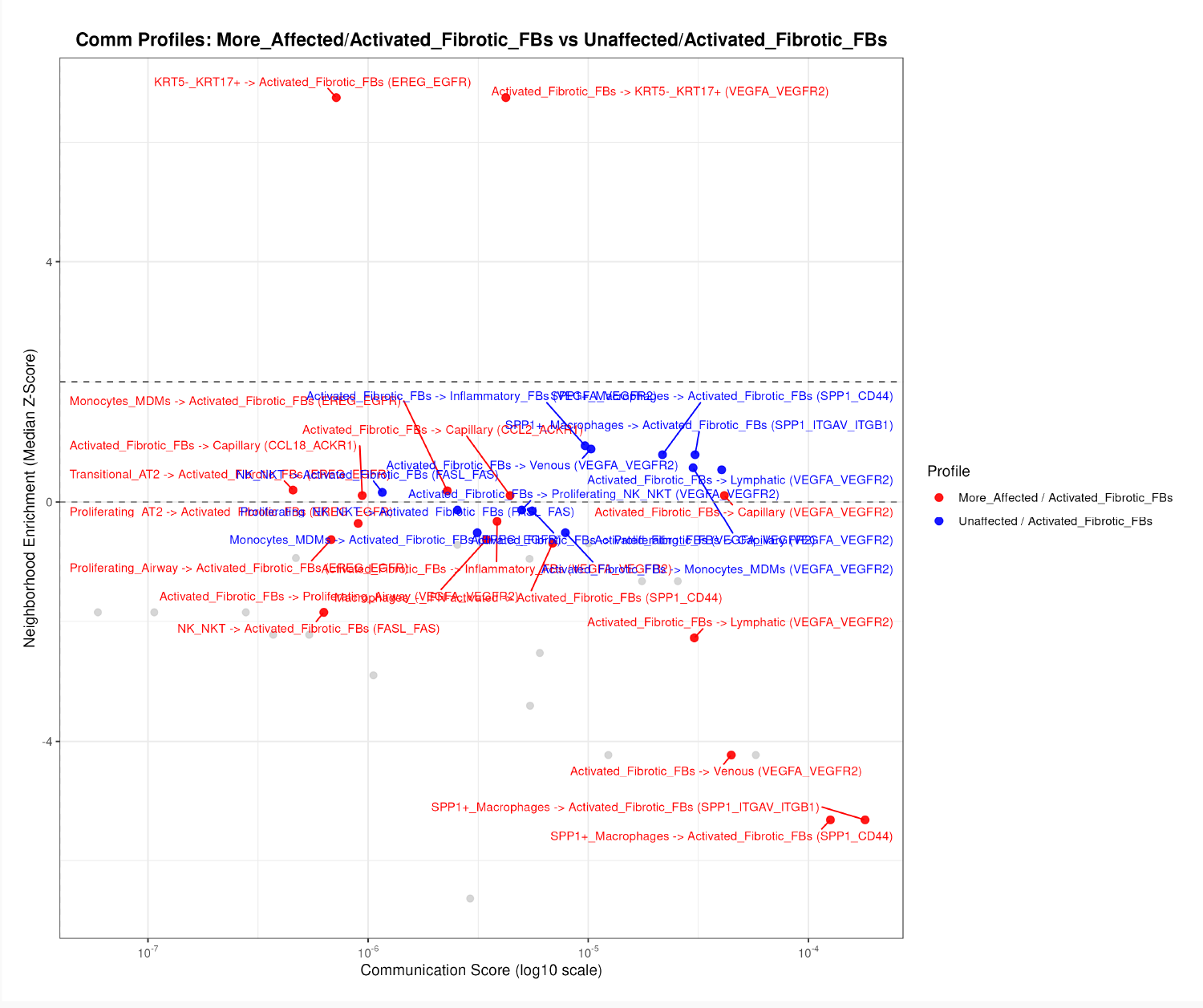
CellChat communication profile for Activated Fibrotic Fibroblasts: More_Affected vs. Unaffected IPF regions. Key gained interactions include SPP1–CD44, VEGFA–VEGFR2, and CCL2–ACKR1 (46 total interactions in More_Affected, 33 unique to the fibrotic state).

##### Step 5 output: iLINCS drug connectivity

The 3,563-gene alveolar fibroblast IPF signature was screened against **2,383 compounds** in the LINCSCP library (reversal mode). The top 15 candidates ranked by anticorrelation score:

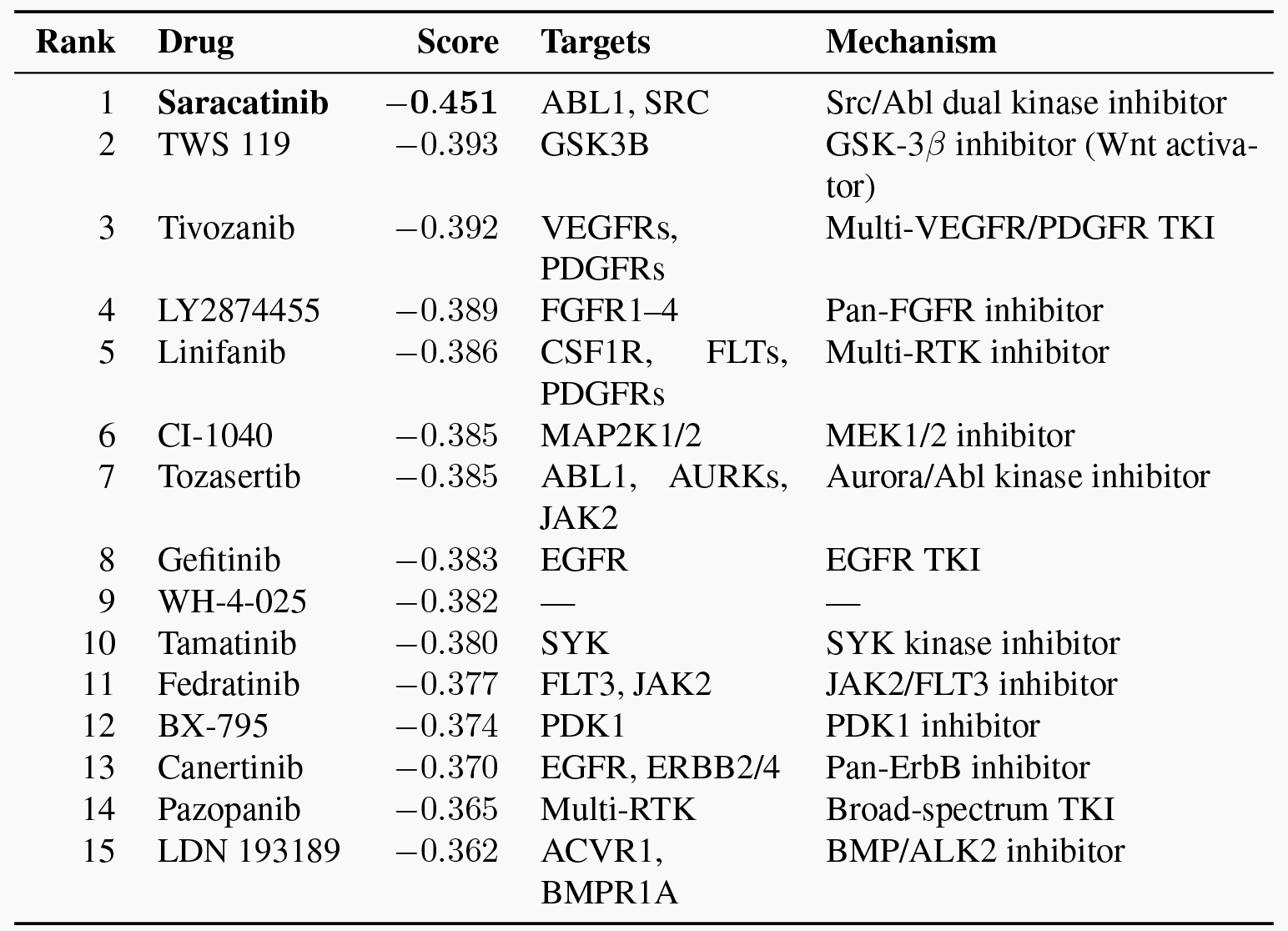

##### Mechanistic convergence

The top hits cluster around Src/Abl kinase, VEGFR/PDGFR signaling, FGFR, MEK/ERK, and JAK2, all pathways directly implicated in the enriched fibroblast program (cell adhesion, migration, ECM remodeling, HIF-1 signaling). Saracatinib was autonomously selected as the priority candidate for downstream DART analysis.

##### Steps 6–7 output: DART cell-type impact analysis

The supervisor agent retrieved Saracatinib’s own iLINCS compound perturbation signature and executed DART, correlating this signature against **475 lung cell-type/disease signatures** across HLCA, BPD_Sun, and HCA_fetal datasets.

**Figure E4:**
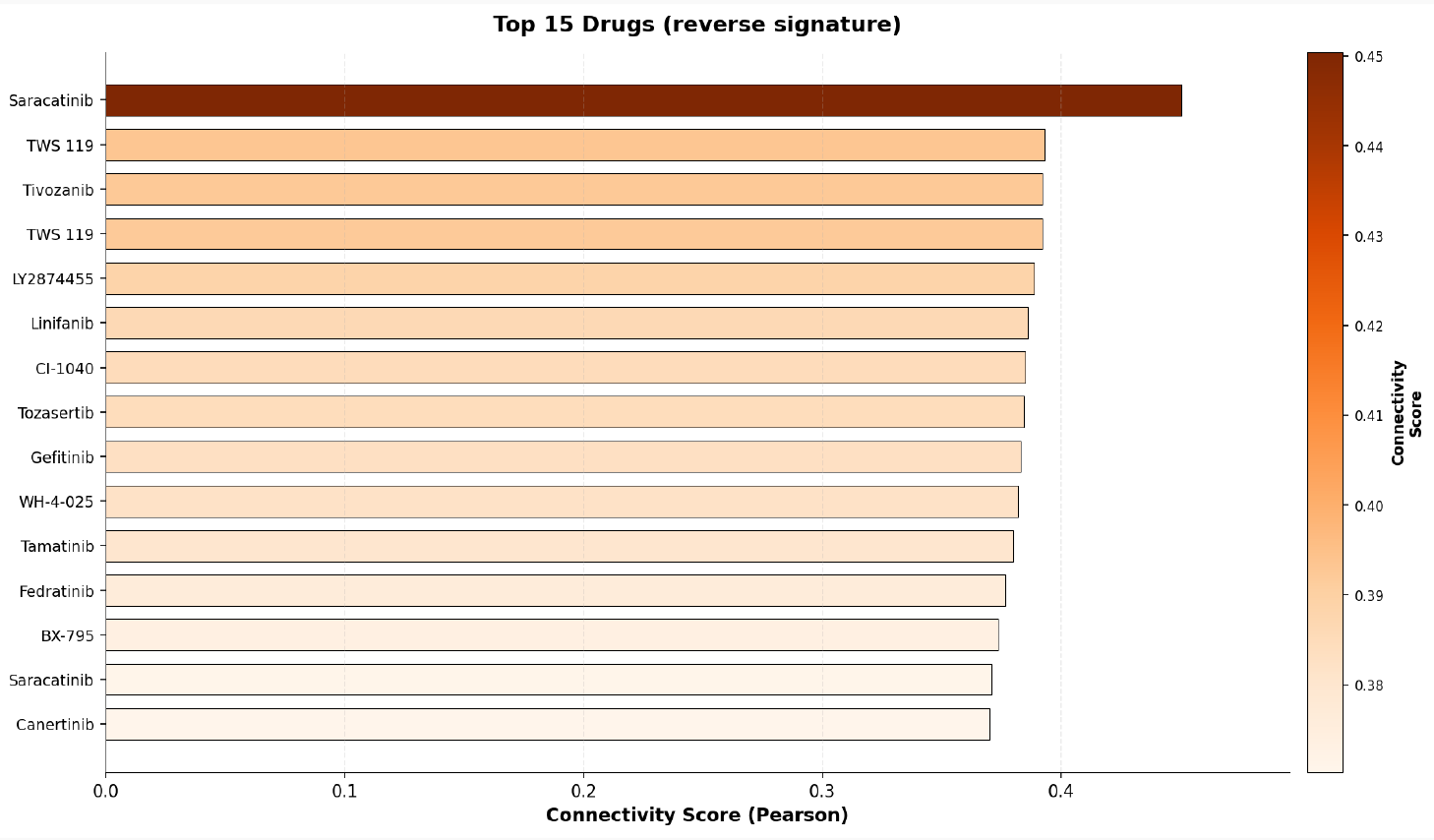
iLINCS drug connectivity screen for the IPF alveolar fibroblast signature (reversal mode, 2,383 compounds). Saracatinib (Src/Abl inhibitor) achieves the highest anticorrelation score (*r* = −0.451), followed by VEGFR/FGFR/MEK inhibitors, consistent with the enriched fibroblast transcriptional program.

**Figure E5:**
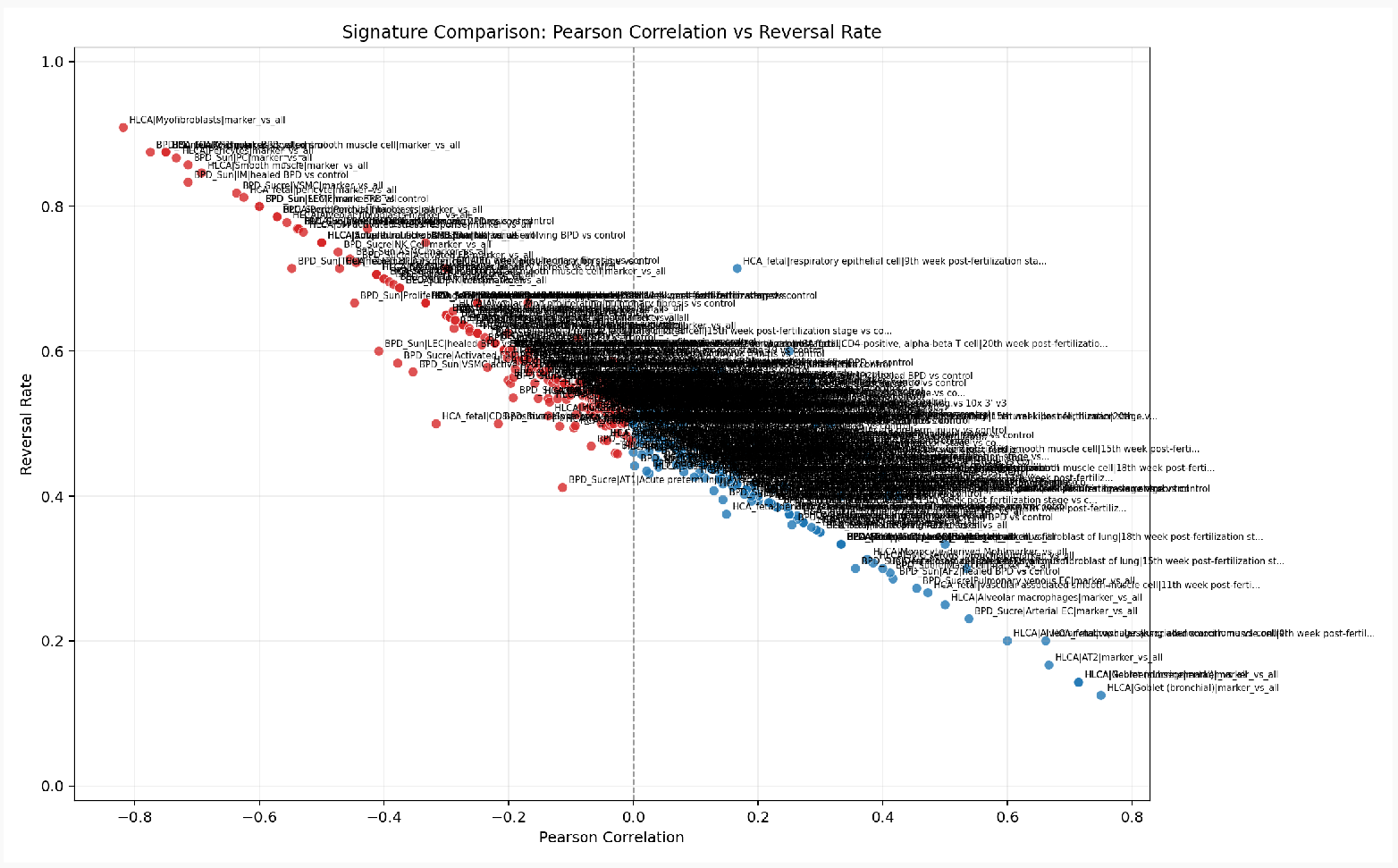
DART cell-type impact analysis for Saracatinib across 475 lung signatures. Negative Pearson *r* (blue) indicates therapeutic reversal of a signature; positive *r* (red) indicates mimicry. Saracatinib achieves strong reversal of myofibroblast (*r* = −0.818), smooth muscle, and pericyte marker programs, with a favorable safety profile in alveolar epithelial cells.

###### Strongest Reversal Signatures (On-Target Therapeutic Benefit)

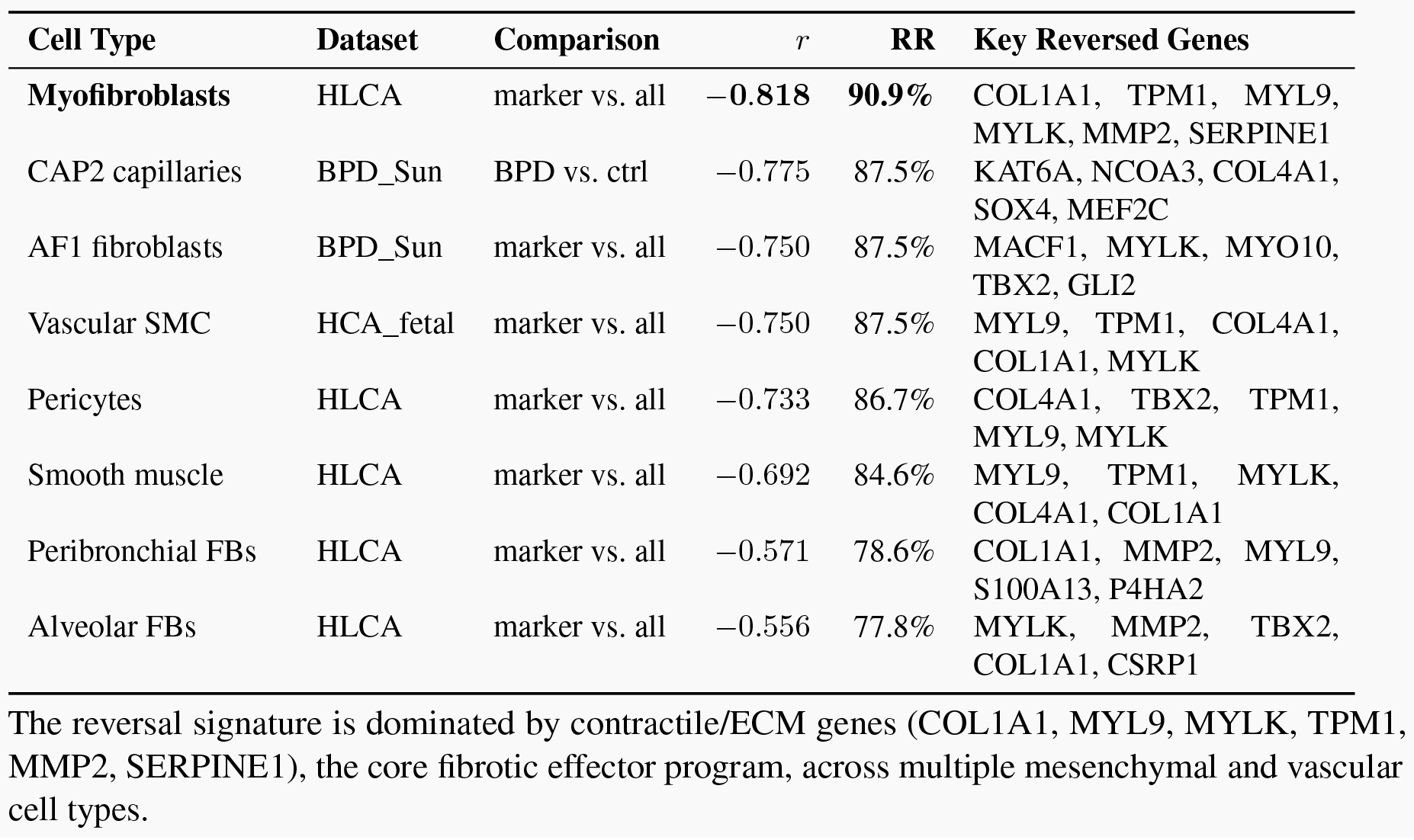

| Cell Type | Dataset | Comparison | $r$ | RR | Key Reversed Genes |
| --- | --- | --- | --- | --- | --- |
| **Myofibroblasts** | HLCA | marker vs. all | **-0.818** | **90.9%** | COL1A1, TPM1, MYL9, MYLK, MMP2, SERPINE1 |
| CAP2 capillaries | BPD_Sun | BPD vs. ctrl | -0.775 | 87.5% | KAT6A, NCOA3, COL4A1, SOX4, MEF2C |
| AF1 fibroblasts | BPD_Sun | marker vs. all | -0.750 | 87.5% | MACF1, MYLK, MYO10, TBX2, GLI2 |
| Vascular SMC | HCA_fetal | marker vs. all | -0.750 | 87.5% | MYL9, TPM1, COL4A1, COL1A1, MYLK |
| Pericytes | HLCA | marker vs. all | -0.733 | 86.7% | COL4A1, TBX2, TPM1, MYL9, MYLK |
| Smooth muscle | HLCA | marker vs. all | -0.692 | 84.6% | MYL9, TPM1, MYLK, COL4A1, COL1A1 |
| Peribronchial FBs | HLCA | marker vs. all | -0.571 | 78.6% | COL1A1, MMP2, MYL9, S100A13, P4HA2 |
| Alveolar FBs | HLCA | marker vs. all | -0.556 | 77.8% | MYLK, MMP2, TBX2, COL1A1, CSRP1 |

The reversal signature is dominated by contractile/ECM genes (COL1A1, MYL9, MYLK, TPM1, MMP2, SERPINE1), the core fibrotic effector program, across multiple mesenchymal and vascular cell types.

###### IPF-Specific Reversal Across Cell Types (HLCA disease signatures)

| Cell Type | Pearson $r$ | Reversal Rate |
| --- | --- | --- |
| EC general capillary | -0.537 | 76.9% |
| Pre-TB secretory | -0.412 | 70.6% |
| EC arterial | -0.300 | 71.4% |
| Alveolar Mph proliferating | -0.289 | 65.5% |
| NK cells | -0.263 | 62.8% |
| CD8 T cells | -0.234 | 57.8% |
| Alveolar fibroblasts | -0.207 | 60.2% |

Saracatinib's reversal effect extends beyond fibroblasts to vascular, immune, and epithelial compartments in IPF, consistent with Src kinase's broad role in cell signaling.

##### Step 8 output: safety profile

DART identified positive correlations between Saracatinib’s perturbation signature and normal cell-type identity programs, indicating potential off-target effects:

| Cell Type | $r$ | Concern |
| --- | --- | --- |
| Goblet (bronchial) | +0.750 | Mimics goblet identity genes (XBP1, SPDEF, MUC1); potential mucus secretion disruption |
| Goblet (subsegmental) | +0.714 | Same mucus program concern |
| Neuroendocrine | +0.714 | Mimics neuroendocrine markers (SNAP25, STMN1); monitor for neuroendocrine effects |
| AT2 cells | +0.667 | Mimics AT2 identity (FGFR2); potential surfactant biology perturbation |
| Alveolar macrophages | +0.500 | Mimics macrophage markers (PPARG, APOE); immune surveillance impact |

##### Interpretation

Saracatinib exhibits strong reversal of fibrotic mesenchymal programs, particularly in myofibroblasts (r = −0.818, 90.9% reversal) and perivascular populations, with the effect extending across vascular and immune compartments, indicating system-level remodeling of the fibrotic niche. The positive correlations with goblet and AT2 cells indicate that Saracatinib’s signature overlaps with normal epithelial identity programs; this does not necessarily indicate toxicity but warrants monitoring of mucociliary function and surfactant biology in preclinical models. LungChat prioritized Saracatinib through iLINCS drug-connectivity screening and then used DART to evaluate its cell-type effects. The compound has been evaluated in the STOP-IPF Phase 1b/2a clinical trial for IPF (NCT04598919), providing external support for the biological relevance of the overall prioritization.

*This entire analysis was completed from a single user prompt in a fresh session with no prior conversation history, demonstrating LungChat’s capacity for autonomous, zero-shot therapeutic discovery with complete provenance.*

### E.2 COPD: Fluticasone Propionate Discovery

#### Supplementary E2. COPD DART Case Study: Cell-Type-Resolved Zero-Shot Therapeutic Discovery

##### User prompt (verbatim)

Identify therapeutic targets and drug repurposing candidates for COPD, separately considering each cell type. Among the top 10 hits for each cell type, which is the most promising therapeutic given prior clinical data for COPD in which cell type. For this drug, show me the top dysregulated genes in that cell population then evaluate its cell-type-specific impact across all lung populations in adults and in utero, to determine on-target efficacy and off-target safety risks.

##### Supervisor decomposition

LungChat’s supervisor agent decomposed this single natural-language prompt into a four-stage pipeline with extensive parallelization across cell types:

1. **Per-cell-type DEG signatures:** Generate COPD vs. control DEGs independently for each of the 14 lung cell types in HLCA
2. **Per-cell-type iLINCS screening:** Run drug connectivity analysis on each cell type’s signature in parallel, producing independent top-10 drug lists per population
3. **Clinical evidence cross-referencing:** Scan all top-10 lists for drugs with documented COPD clinical trial history; select the most clinically validated candidate
4. **DART cell-type impact:** For the selected drug, generate a volcano plot of DEGs in its primary cell type; then execute DART across adult (HLCA) and fetal (HCA_fetal) lung populations

Unlike the IPF case study, this workflow independently screens **14 cell types**, enabling cell-type-resolved drug discovery and clinical evidence cross-validation across populations.

##### Complete execution trace

**Complete execution trace.**
| Step | Agent | Tool(s) | Operation |
| --- | --- | --- | --- |
| 1 | Supervisor | decompose | Parses query; identifies 14 cell types with COPD samples in HLCA; plans parallel iLINCS dispatch |
| 2 | Single-Cell | generate_stats ($\times 14$) | Differential expression for each of 14 cell types (COPD vs. control, HLCA). 8 cell types yield signatures with sufficient gene counts for iLINCS |
| 3 | Research | drug_connectivity_tool ($\times 8$, batched) | iLINCS drug connectivity analysis on each cell type's DEG signature independently (reversal mode, LINCSCP). Returns per-cell-type top-10 ranked drug lists |
| 4 | Research | literature_search_tool,<br>clinical_trials_search_tool | Cross-references all top-10 hits against COPD clinical trial databases. Identifies Fluticasone Propionate (>200 trials, Phase 4 approved) in DC2 and Non-classical Monocyte lists |
| 5 | Single-Cell | plot_differential_expression_volcano | Volcano plot of 521 DEGs in DC2 cells (COPD vs. control) |
| 6 | Supervisor | query_pseudobulk_expression_data_tool (drug_cell_type_impact) | Executes DART: correlates Fluticasone Propionate iLINCS signature (978 genes) against 252 lung signatures across HLCA (adult) and HCA_fetal (in utero) datasets |
| 7 | Supervisor | Synthesis | Aggregates all outputs into integrated therapeutic report with on-target efficacy and fetal off-target safety profiling |

### Step 2–3 output: per-cell-type drug screening

Of 14 cell types with COPD DEG signatures, 8 yielded sufficient gene counts for iLINCS analysis. Independent screening produced cell-type-resolved drug rankings:

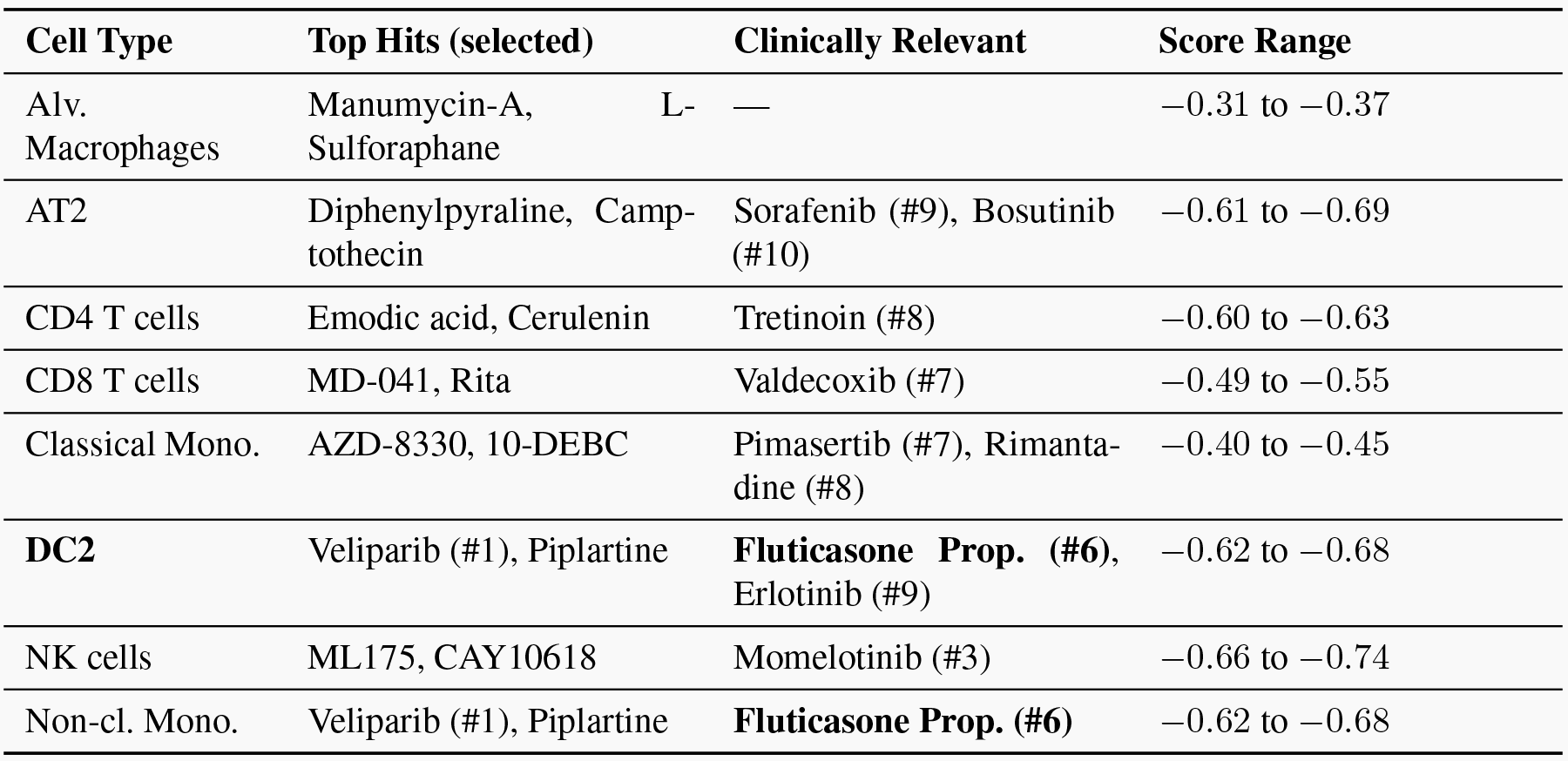

### Step 4 output: clinical evidence cross-referencing

Scanning all 80 candidate drugs (top 10 × 8 cell types) against COPD clinical trial databases:

| Drug | Cell Types (rank) | COPD Clinical Evidence |
| --- | --- | --- |
| **Fluticasone Propionate** | DC2 (#6), Non-cl. Mono. (#6) | **Approved.** >200 COPD trials. Phase 4 completed. |
| Tretinoin | CD4 T cells (#8) | Phase 2 only (emphysema subtype). Not approved. |
| Rimantadine | Classical Mono. (#8) | One trial (viral prophylaxis context only). |
| Others | Various | No COPD trials identified. |

**Fluticasone Propionate** was selected as the priority candidate: the only drug in any top-10 list with Phase 4 COPD approval, emerging independently from both DC2 and Non-classical Monocyte signatures (*p* = 3.4 × 10^−5^).

### Step 5 output: DC2 differential expression (COPD vs. control)

Analysis of DC2 (type 2 dendritic) cells identified **521 DEGs** (COPD vs. control). The top dysregulated genes reveal a shift toward a mature, antigen-presenting phenotype with loss of innate alarm signaling:

| Dir. | Gene | LogFC | FDR | Functional Role |
| --- | --- | --- | --- | --- |
| ↑ | CD207 | +2.49 | $9.95 \times 10^{-5}$ | DC maturation / antigen capture |
| ↑ | PKP2 | +2.16 | $6.22 \times 10^{-7}$ | Desmosomal signaling |
| ↑ | MT2A | +1.77 | $1.13 \times 10^{-5}$ | Metallothionein, oxidative stress |
| ↑ | F2RL2 | +1.69 | $2.86 \times 10^{-4}$ | Protease-activated receptor |
| ↑ | CXCL9 | +1.64 | $4.13 \times 10^{-4}$ | T cell recruitment chemokine |
| ↓ | S100A9 | -1.74 | $4.42 \times 10^{-5}$ | Alarmin / innate immune activation |
| ↓ | CXCL8 | -1.73 | $1.50 \times 10^{-4}$ | Neutrophil recruitment (IL-8) |
| ↓ | MMP14 | -1.55 | $5.58 \times 10^{-4}$ | Matrix metalloprotease |
| ↓ | SIRPA | -1.25 | $2.64 \times 10^{-6}$ | Phagocytosis checkpoint |
| ↓ | TNFAIP8 | -1.20 | $6.57 \times 10^{-6}$ | NF-κB / apoptosis regulator |
| ↓ | HIF1A | -1.05 | $1.28 \times 10^{-5}$ | Hypoxia response |
| ↓ | BCL2A1 | -1.03 | $3.29 \times 10^{-4}$ | Anti-apoptotic |

**Figure E6:**
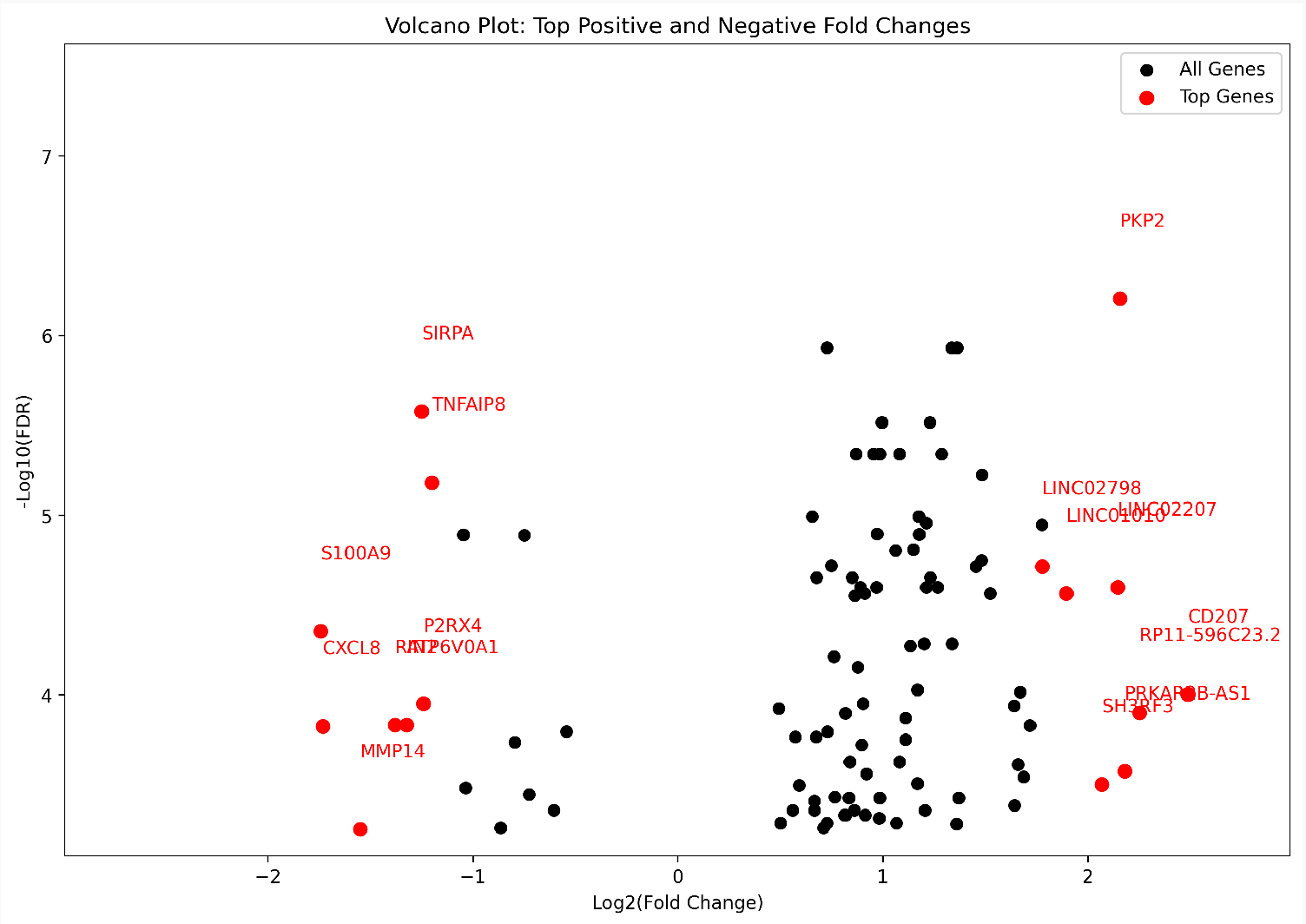
Volcano plot of differential gene expression in COPD vs. control DC2 cells (521 DEGs). Upregulated genes (CD207, CXCL9, PKP2) indicate enhanced antigen presentation; downregulated genes (S100A9, CXCL8, MMP14, HIF1A) indicate loss of innate alarm signaling and hypoxia response.

### Step 6 output: DART cell-type impact analysis

The Fluticasone Propionate iLINCS perturbation signature (**978 genes**) was correlated against **252 lung signatures** across HLCA (adult) and HCA_fetal (in utero) datasets. A 100-gene subset was used for impact analysis.

**Figure E7:**
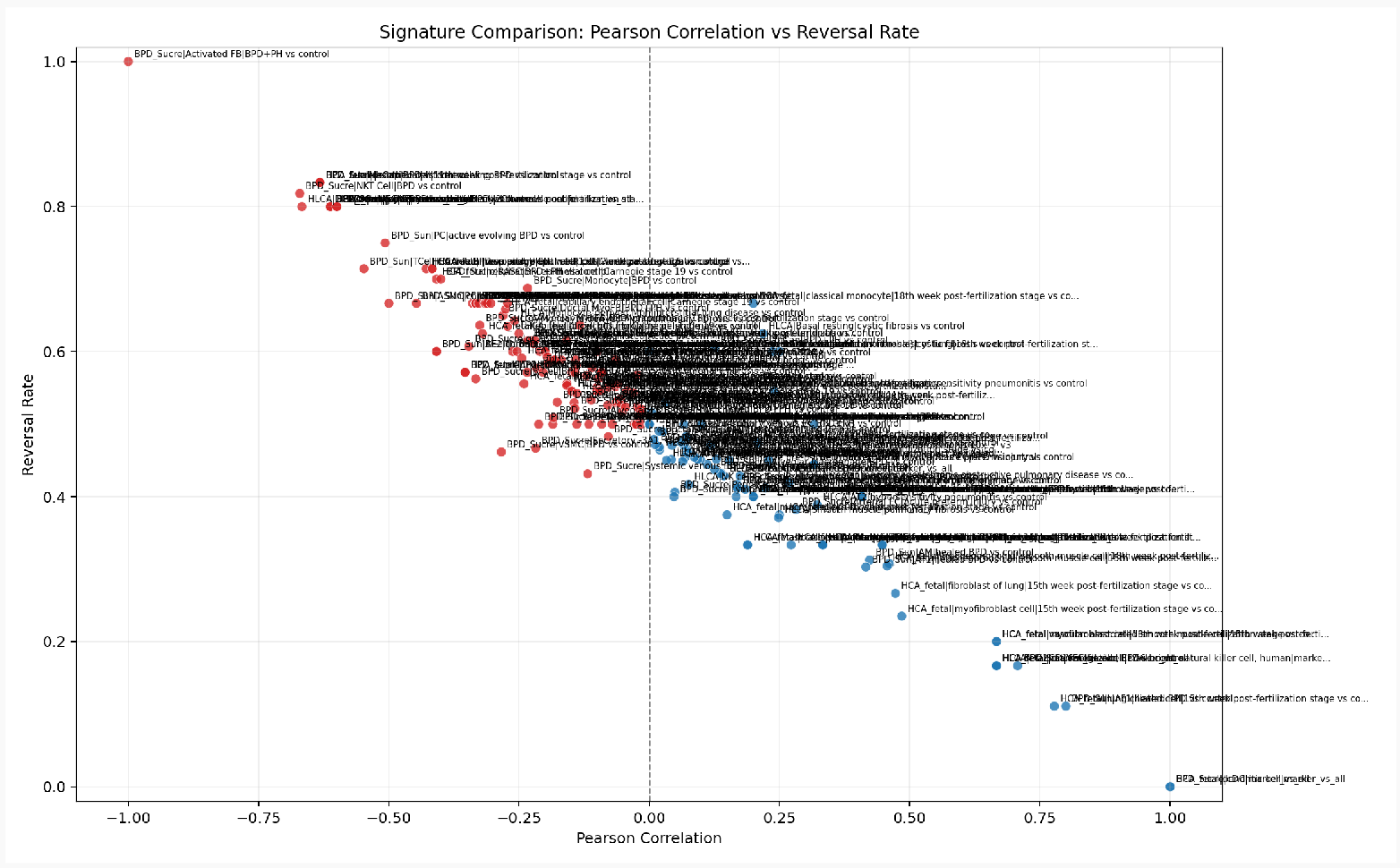
DART cell-type impact analysis for Fluticasone Propionate across 252 lung signatures (HLCA adult + HCA_fetal). Negative Pearson *r* (blue) indicates therapeutic reversal; positive *r* (red) indicates signature amplification. DC2 COPD signature shows strong reversal (*r* = −0.667, 80% RR), while fetal dendritic and ciliated cell developmental programs show positive correlation, indicating potential off-target effects during gestation.

#### On-Target Efficacy (Adult HLCA, COPD Signatures)

| Cell Type | Pearson $r$ | RR | Overlap | Key Reversed Genes |
| --- | --- | --- | --- | --- |
| DC2 (COPD vs. ctrl) | -0.667 | 80% | 5 | JUN↓, USP7↓, CCNH↓, MEF2C↓ |

The DC2 COPD signature is the only adult HLCA COPD signature with sufficient overlap, and it shows strong anticorrelation (*r* = −0.667, reversal rate 80%), confirming Fluticasone’s predicted on-target activity in the cell type where it was identified.

#### Off-Target Safety (Fetal HCA_fetal, Developmental Programs)

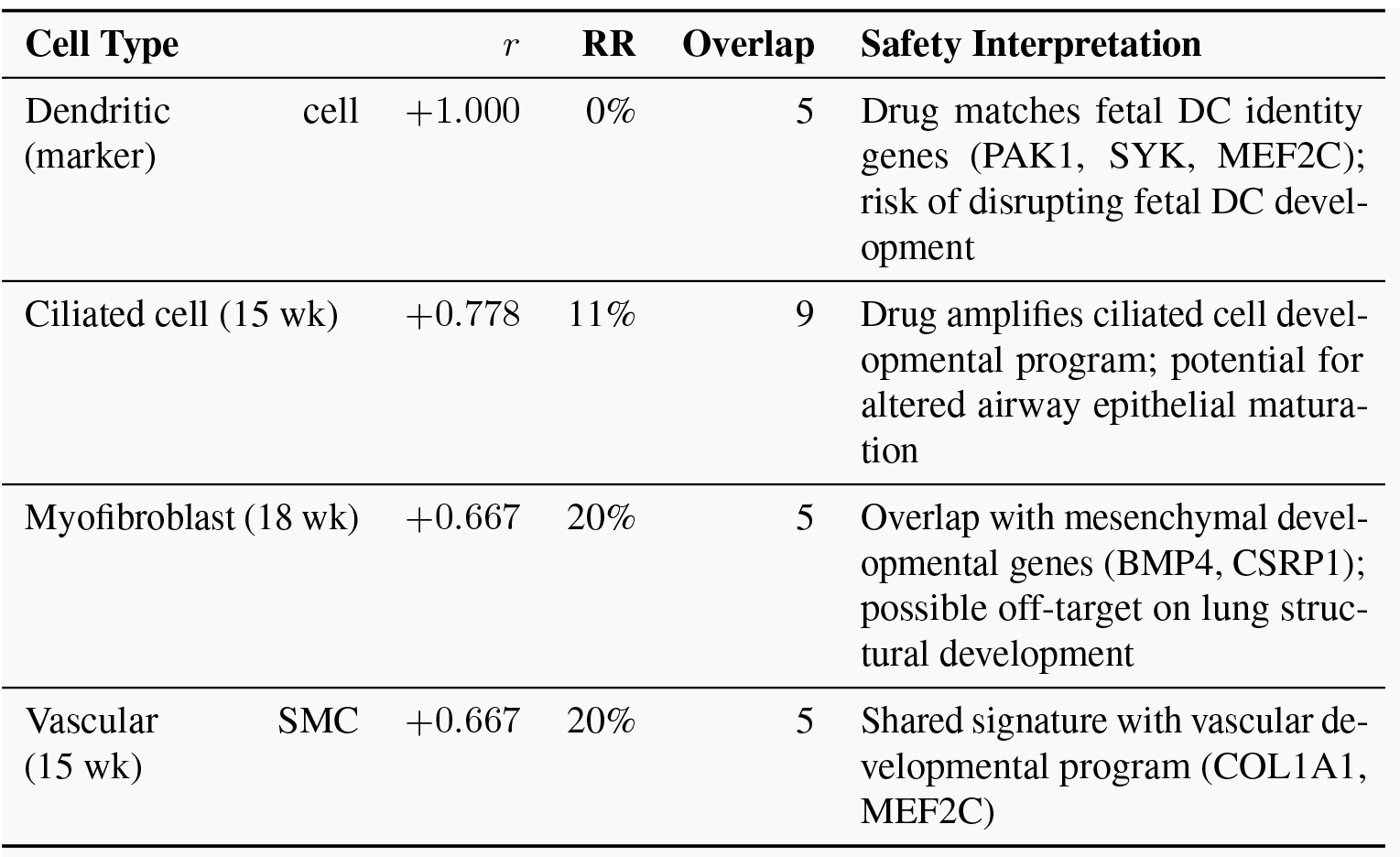

### Interpretation

Fluticasone Propionate demonstrates strong reversal of the COPD-associated transcriptional program in DC2 dendritic cells (*r* = −0.667, 80% reversal rate), the primary cell type where it was computationally identified. The reversed genes (JUN, MEF2C, CCNH, USP7) are consistent with glucocorticoid receptor (NR3C1)-mediated suppression of inflammatory signaling, mechanistically linking the drug’s known anti-inflammatory activity to the COPD DC2 disease program.

The fetal safety analysis reveals a clinically interpretable risk: Fluticasone’s signature positively correlates with normal developmental programs in dendritic cells (*r* = +1.0), ciliated epithelial cells (*r* = +0.778), myofibroblasts (*r* = +0.667), and vascular smooth muscle (*r* = +0.667) at 15–18 weeks gestation. This transcriptomic signal is consistent with the need for caution and pregnancy/lactation risk assessment in current drug-labeling frameworks, providing cell-type-resolved molecular evidence for potential effects during fetal airway and mesenchymal development.

The finding that an approved, Phase 4-validated COPD corticosteroid is autonomously identified as the top candidate from cell-type-resolved computational screening, with mechanistically coherent on-target and off-target profiles, provides strong external validation of the DART method’s capacity for clinically meaningful drug repurposing. The overlap counts are small (5–9 genes per comparison), so these correlations should be interpreted as directional signals rather than definitive predictions.

*This entire analysis, including per-cell-type drug screening across 14 populations, clinical evidence cross-referencing, DC2 differential expression, and dual-dataset (adult + fetal) DART safety profiling,was completed from a single user prompt in a fresh session with no prior conversation history.*

## F Architecture ablation details

This appendix provides the full specification and detailed results for the architecture ablation described in Section 2.2.

### F.1 Ablation configuration details

Table F1 details the four architecture configurations evaluated. The key design principle: flat-agent baselines preserve all domain knowledge from specialist prompts, ensuring that observed differences reflect architectural choices rather than information asymmetry.

**Table F1:** Ablation arm specifications. All configurations use the same foundation models, datasets, and evaluation rubrics.

| Config | Agent(s) | Prompt | Tool Access | What It Isolates |
| --- | --- | --- | --- | --- |
| MA+Sel (production) | 5 agents (Supervisor + 4 specialists) | Specialized per agent | Supervisor routing + hybrid selector | Full system baseline |
| MA−Sel | 5 agents (Supervisor + 4 specialists) | Specialized per agent | Supervisor routing, no selector | Selector contribution |
| Flat+Sel | 1 agent | Merged flat prompt | All 51 tools + hybrid selector | Value of hierarchy when tool narrowing retained |
| Flat−Sel | 1 agent | Merged flat prompt | All 51 tools, no selector | Hierarchy + tool narrowing combined |

### F.2 Context engineering rationale

The multi-agent architecture is motivated by a context-engineering principle: each specialist agent operates in a scoped context window containing only its domain-relevant tools and instructions, preventing cross-domain parameter leakage and instruction dilution. Token efficiency is a measurable consequence of this design (cumulative prompt tokens scale sub-linearly with task complexity), but the primary benefit is preserved reasoning quality under increasing tool and domain complexity.

Table F2 summarizes the specific context problems addressed by the hierarchical design.

**Table F2:**
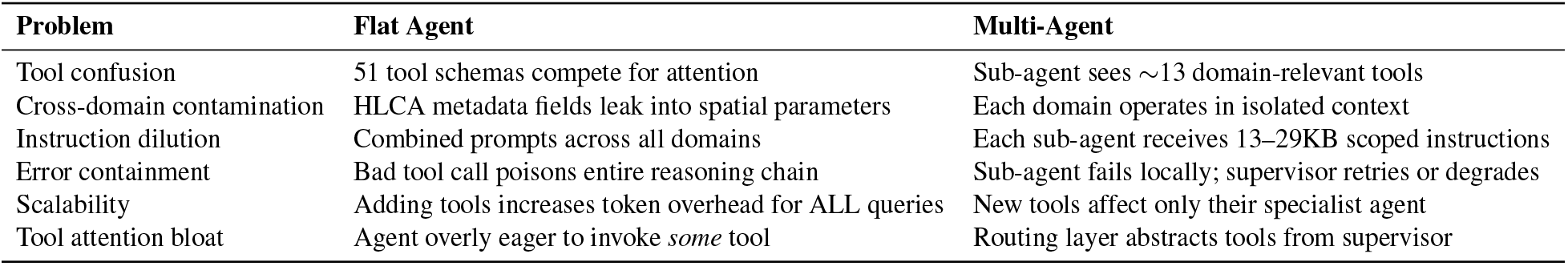
Context problems addressed by multi-agent hierarchy vs. flat architecture.

| Problem | Flat Agent | Multi-Agent |
| --- | --- | --- |
| Tool confusion | 51 tool schemas compete for attention | Sub-agent sees ~13 domain-relevant tools |
| Cross-domain contamination | HLCA metadata fields leak into spatial parameters | Each domain operates in isolated context |
| Instruction dilution | Combined prompts across all domains | Each sub-agent receives 13–29KB scoped instructions |
| Error containment | Bad tool call poisons entire reasoning chain | Sub-agent fails locally; supervisor retries or degrades |
| Scalability | Adding tools increases token overhead for ALL queries | New tools affect only their specialist agent |
| Tool attention bloat | Agent overly eager to invoke *some* tool | Routing layer abstracts tools from supervisor |

The context-partitioning principles underlying our design align with several recent findings. Liu et al. [15] demonstrated systematic “lost in the middle” degradation where LLM accuracy drops for mid-context information, a phenomenon exacerbated by monolithic context accumulation. Anthropic [2] identifies subagent delegation and context compaction as primary mitigations for context degradation in long-running agents. Zhang et al. [30] showed that multi-agent collaboration through scoped worker contexts outperforms single-agent systems by up to 10% on long-context tasks.

### F.3 Full single-step ablation results

Table F3 provides the complete per-metric breakdown for all 50 single-step queries across 4 ablation modes (*n*=3 repetitions each, 600 total evaluations).

**Table F3:** Full single-step ablation results (50 queries × 3 reps × 4 modes = 600 evaluations). **Bold** indicates best per metric. Quality: higher is better. Efficiency: lower is better.

|  | Metric | MA+Sel | MA−Sel | Flat+Sel | Flat−Sel |
| --- | --- | --- | --- | --- | --- |
| Quality | LLM Judge | $0.893 \pm 0.31$ | $0.927 \pm 0.26$ | $0.947 \pm 0.23$ | **$0.980 \pm 0.14$** |
| | Tool Selection | $0.920 \pm 0.27$ | $0.933 \pm 0.25$ | $0.927 \pm 0.24$ | **$0.980 \pm 0.14$** |
| | Config Match | $0.927 \pm 0.26$ | $0.940 \pm 0.24$ | $0.913 \pm 0.28$ | **$0.947 \pm 0.23$** |
| | Items Coverage | $0.891 \pm 0.27$ | $0.919 \pm 0.23$ | **$0.927 \pm 0.22$** | $0.924 \pm 0.21$ |
| | Execution Success | $0.973 \pm 0.16$ | **$0.980 \pm 0.14$** | $0.973 \pm 0.16$ | $0.973 \pm 0.16$ |
| | Metadata Consistency | **$1.000 \pm 0.00$** | $0.973 \pm 0.16$ | $0.973 \pm 0.16$ | **$1.000 \pm 0.00$** |
| Efficiency | Mean Latency (s) | 50.2 | 56.7 | 46.3 | **39.5** |
|  | Mean Tokens (K) | **197** | 212 | 330 | 401 |
|  | Mean Tool Calls | **2.05** | 2.18 | 3.79 | 3.37 |

#### Observations

1. **Quality**: Flat−Sel achieves the highest LLM Judge score (0.980) and Tool Selection accuracy (0.980), but the absolute gap to the production configuration (MA+Sel) is ≤0.09 across all metrics.
2. **Token efficiency**: MA+Sel consumes ∼2× fewer tokens than Flat−Sel (197K vs. 401K), reflecting context partitioning across specialist agents.
3. **Latency**: Flat−Sel is fastest (39.5s) because it avoids supervisor→sub-agent routing overhead, while MA+Sel adds ∼10s for delegation.
4. **Tool calls**: MA+Sel averages 2.05 tool calls vs. Flat−Sel’s 3.37, because flat agents attempt more exploratory tool calls without hierarchical guidance.
5. **Metadata consistency**: MA+Sel achieves perfect metadata consistency (1.000), suggesting that scoped contexts improve parameter grounding for dataset-specific metadata fields.

### F.4 Security and grounded abstention results

#### Security (13 queries × 3 reps × 4 modes = 156 evaluations)

All four architectures achieved 100% prevention of explicit tool execution and secret leakage under prompt injection, demonstrating that core supervisor prompt engineering and sandbox boundaries are robustly engineered regardless of tool configuration. MA+Sel showed slightly lower refusal coherence (82.1% vs. 92.3%) because the selector mechanism occasionally attempts to legitimately route complex adversarial language before recognizing hostile intent.

#### Grounded abstention (17 queries × 3 reps × 4 modes = 204 evaluations)

This benchmark provides the strongest architectural validation. When presented with out-of-scope queries, flat architectures (whose context windows expose all 51 tool schemas) become overly eager to invoke *some* tool (54.9% disallowed-tool blocking vs. 90.2% for MA+Sel). This +35% absolute improvement directly validates the context-engineering hypothesis: abstracting domain tools behind a routing layer forces the top-level agent to reason about intent before invoking capabilities, yielding superior grounded refusal. MA+Sel also produces fewer invalid output artifacts (86.3% clean vs. 70.6% for Flat−Sel), indicating that context isolation reduces spurious output generation.

### F.5 Multi-tool orchestration results

Based on the monotonic cost-quality ordering in the 4-arm single-step ablation (MA+Sel: best cost, Flat−Sel: best accuracy), the multi-tool and decomposition-stress benchmarks were evaluated on the two boundary configurations: MA+Sel (production) and Flat−Sel.

#### Quality (10 queries × 3 reps × 2 modes = 60 evaluations)

Table F4 summarizes multi-tool orchestration results. Both architectures achieve identical strict pass rates (66.7%) and comparable mean rubric scores (MA+Sel: 3.10/4, Flat−Sel: 3.23/4). Per-question analysis reveals task-dependent architecture preference rather than a global advantage: MA+Sel wins on queries requiring gene regulatory network generation with literature search or multi-dataset cell frequency analysis (MT03, MT04, MT06, MT08), where the supervisor’s decomposition into parallel sub-agent tasks produces more coherent synthesis. Flat−Sel wins on queries with broader tool orchestration (MT01, MT05, MT09, MT10), where direct access to all tools avoids routing overhead.

**Table F4:** Multi-tool orchestration ablation (10 queries × 3 reps × 2 modes). Quality: higher is better. Efficiency: lower is better. **Bold** indicates best per metric.

|  | **Metric** | **MA+Sel** | **Flat–Sel** |
| --- | --- | --- | --- |
| Quality | Execution Success | 0.967 ± 0.18 | **1.000** ± 0.00 |
| | Mean Rubric Score (0–4) | 3.10 ± 1.51 | **3.23** ± 1.36 |
|  | Strict Pass Rate (%) | 66.7 | 66.7 |
|  | Task Coverage (%) | 80.0 | **83.3** |
|  | Trace Fidelity (%) | 76.7 | **80.0** |
|  | Cross-Domain Synthesis (%) | 73.3 | **76.7** |
| Efficiency | Mean Latency (s) | 303.5 | **151.7** |
|  | Mean Tokens (K) | **1,477** | 1,540 |
|  | Mean Tool Calls | **12.0** | 18.6 |
|  | Hard Failure Rate (%) | 10.0 | **6.7** |

#### Efficiency

Flat−Sel is 2× faster (152s vs. 304s) due to the absence of supervisor→sub-agent delegation overhead. However, total token consumption is nearly identical (∼1.5M per query), indicating the latency difference arises from sequential API calls in the delegation chain, not from additional computation. Flat−Sel uses 55% more tool calls (18.6 vs. 12.0), compensating for the lack of hierarchical routing intelligence with broader exploratory tool invocation. The cost crossover relative to single-step queries is notable: on multi-tool queries, Flat−Sel is modestly cheaper, because each sub-agent invocation in the hierarchical architecture carries context setup overhead.

### F.6 Decomposition stress results

#### Quality (10 queries × 3 reps × 2 modes = 60 evaluations)

Table F5 summarizes decomposition-stress results. Both architectures achieve near-ceiling quality: *>*83% strict pass rates with zero hard failures and mean rubric scores *>*5.3/6. This is the highest-quality benchmark in the evaluation suite, demonstrating that LungChat’s analytical quality on complex decomposition tasks comes from its domain engineering (curated tools, pre-computed warehouse, expert prompts) rather than from the multi-agent hierarchy.

**Table F5:**
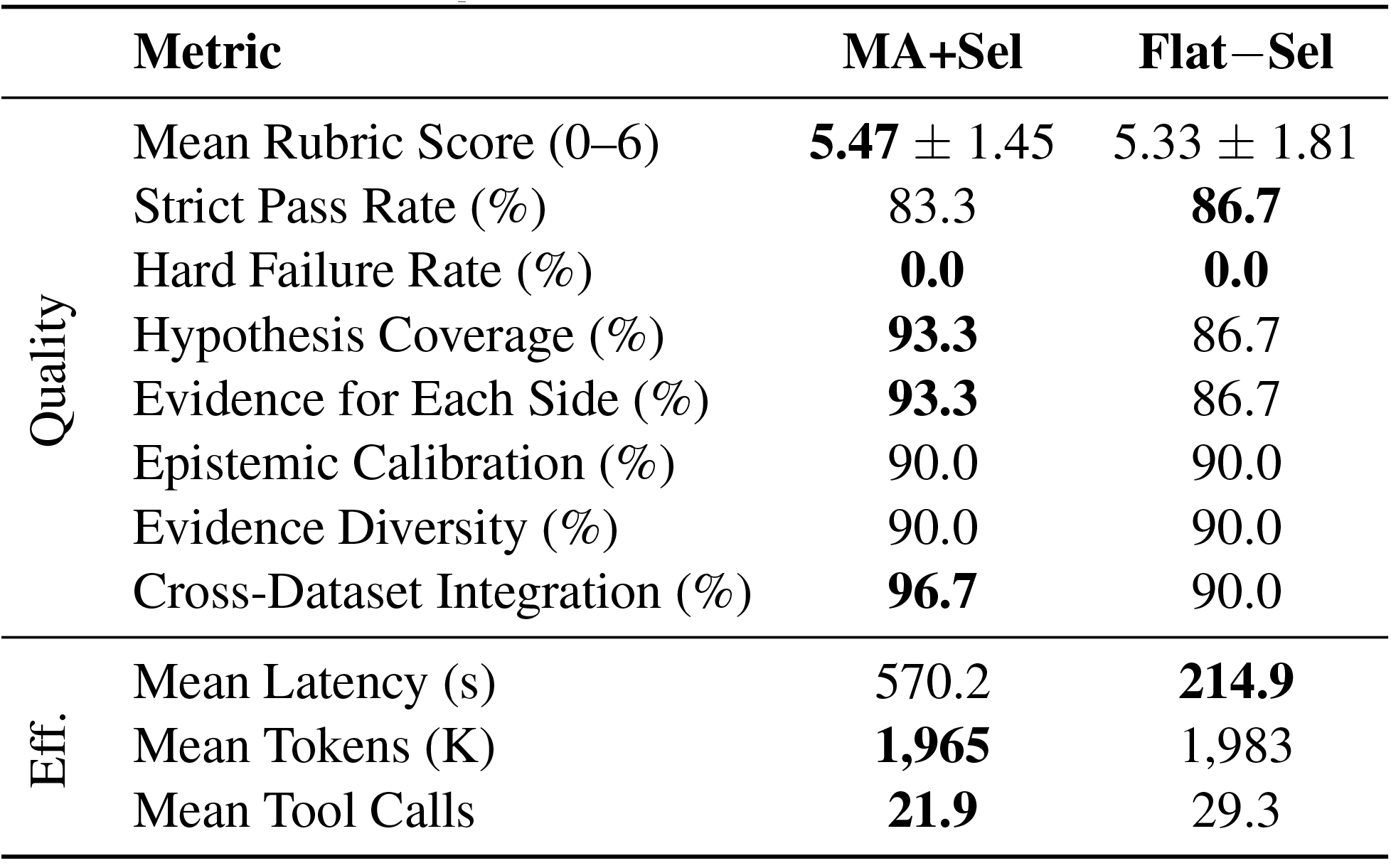
Decomposition stress ablation (10 queries × 3 reps × 2 modes). Quality: higher is better. Efficiency: lower is better. **Bold** indicates best per metric.

|  | **Metric** | **MA+Sel** | **Flat–Sel** |
| --- | --- | --- | --- |
| Quality | Mean Rubric Score (0–6) | **5.47** ± 1.45 | 5.33 ± 1.81 |
|  | Strict Pass Rate (%) | 83.3 | **86.7** |
|  | Hard Failure Rate (%) | **0.0** | **0.0** |
|  | Hypothesis Coverage (%) | **93.3** | 86.7 |
|  | Evidence for Each Side (%) | **93.3** | 86.7 |
|  | Epistemic Calibration (%) | 90.0 | 90.0 |
|  | Evidence Diversity (%) | 90.0 | 90.0 |
|  | Cross-Dataset Integration (%) | **96.7** | 90.0 |
| Eff. | Mean Latency (s) | 570.2 | **214.9** |
|  | Mean Tokens (K) | **1,965** | 1,983 |
|  | Mean Tool Calls | **21.9** | 29.3 |

#### Mechanistic observations

MA+Sel shows a modest edge on criteria that specifically reward structured decomposition: hypothesis coverage (+6.6pp), evidence for each side (+6.6pp), and cross-dataset integration (+6.7pp). These criteria reward the supervisor’s ability to dispatch independent analyses to specialist agents before synthesis. Flat−Sel matches or exceeds on mechanistic depth (90.0% vs. 83.3%), suggesting that direct tool access without routing intermediation can yield deeper single-branch reasoning. Flat−Sel is 2.7× faster (215s vs. 570s) with dramatically lower variance, making it more predictable for complex queries.

### F.7 Combined evaluation summary

Table F6 provides a consolidated view across all five evaluation tracks (100 unique questions, 1,080 total evaluations). No single configuration dominates all dimensions.

**Table F6:** Combined evaluation summary across all five tracks. **Bold** indicates the better configuration per metric. Quality metrics are reported as percentages; token counts are in thousands.

| Track (N) | Key Metric | MA+Sel | Flat–Sel | Winner |
| --- | --- | --- | --- | --- |
| Single-Step (600) | LLM Judge (%) | 89.3 | **98.0** | Flat (−8.7pp) |
| Single-Step (600) | Mean Tokens (K) | **197** | 401 | MA (2× fewer) |
| Security (156) | No Leak / No Exec (%) | **100.0** | **100.0** | Tie |
| Security (156) | Mean Tokens (K) | **38** | 89 | MA (2.3× fewer) |
| Grounded Abstention (204) | No Disallowed Tools (%) | **90.2** | 54.9 | **MA (+35.3pp)** |
| Grounded Abstention (204) | Mean Tokens (K) | **131** | 340 | MA (2.6× fewer) |
| Multi-Tool (60) | Mean Rubric (0–4) | 3.10 | **3.23** | ≈Tie (−0.13) |
| Multi-Tool (60) | Mean Tokens (K) | **1,477** | 1,540 | ≈Tie |
| Decomp. Stress (60) | Mean Rubric (0–6) | **5.47** | 5.33 | ≈Tie (+0.14) |
| Decomp. Stress (60) | Mean Tokens (K) | **1,965** | 1,983 | ≈Tie |

The production multi-agent architecture (MA+Sel) achieves the best profile across the Pareto frontier: 89% single-step accuracy (vs. 98% for Flat−Sel), 90% disallowed-tool blocking (vs. 55% for Flat−Sel), comparable multi-tool and decomposition-stress reasoning, and 2× token efficiency on the dominant single-step query class. The flat-agent baseline demonstrates that LungChat’s domain knowledge, not its architecture, is the primary driver of analytical quality, while the multi-agent hierarchy provides the safety, cost, and scalability properties required for production deployment.

#### F.8 Discussion and limitations

Across all five evaluation tracks (100 questions, 1,080 evaluations), analytical quality is architecture-neutral: both multi-agent and flat-agent configurations achieve comparable accuracy on single-step, multi-tool, and decomposition-stress queries. The multi-agent hierarchy’s measurable contribution is on dimensions that accuracy-only benchmarks do not capture: a +35pp improvement in grounded abstention (preventing the system from invoking biological tools on out-of-scope queries), 2× token-cost reduction on the dominant single-step query class, and structural guarantees for error containment and extensibility.

The current evaluation focuses on single-turn queries. While the stateless sub-agent design provides structural advantages for multi-turn session continuity, where each sub-agent invocation operates in a fresh context independent of prior turns, we have not empirically evaluated this property across long sessions. At https://chat.lungmap.net, researchers routinely conduct sessions spanning 10–20 queries; in a monolithic architecture, accumulated context would approach 4M+ tokens after 10 turns, while the multi-agent supervisor carries compact summaries (∼100K tokens) with each sub-agent receiving a fresh context. Quantifying quality degradation over extended sessions is an important direction for future work.

A natural next step is to distill the current LLM-based tool reranker into a lightweight learned reranker trained on successful traces, while retaining LLM fallback for low-confidence or out-of-distribution cases.

## Notes

### Competing Interest Statement

The authors have declared no competing interest.

